# Adaptation shapes the representational geometry in mouse V1 to efficiently encode the environment

**DOI:** 10.1101/2024.12.11.628035

**Authors:** Mario Dipoppa, Ramon Nogueira, Stéphane Bugeon, Yoni Friedman, Charu B. Reddy, Kenneth D. Harris, Dario L. Ringach, Kenneth D. Miller, Matteo Carandini, Stefano Fusi

**Affiliations:** Department of Neurobiology, University of California, Los Angeles, CA, USA; Department of Computational Medicine, University of California, Los Angeles, CA, USA; Center for Theoretical Neuroscience, Zuckerman Institute for Brain Mind and Behavior, Columbia University, NY, USA; Institute of Neurology, University College London, UK; Grossman Center for Quantitative Biology and Human Behavior, University of Chicago, Chicago, IL, USA; Department of Neurobiology, University of Chicago, Chicago, IL, USA; Massachusetts Institute of Technology, MA, USA; Department of Psychology, University of California, Los Angeles, CA, USA; Kavli Institute for Brain Science, Columbia University, NY, USA; Institute of Ophthalmology, University College London, UK

**Author notes:** contributed equally (Ramon Nogueira, Stéphane Bugeon). co-senior authors (Mario Dipoppa, Matteo Carandini, Stefano Fusi).

## Abstract

Sensory adaptation dynamically changes neural responses as a function of previous stimuli, profoundly impacting perception. The response changes induced by adaptation have been characterized in detail in individual neurons and at the population level after averaging across trials. However, it is not clear how adaptation modifies the aspects of the representations that relate more directly to the ability to perceive stimuli, such as their geometry and the noise structure in individual trials. To address this question, we recorded from a population of neurons in the mouse visual cortex and presented one stimulus (an oriented grating) more frequently than the others. We then analyzed these data in terms of representational geometry and studied the ability of a linear decoder to discriminate between similar visual stimuli based on the single-trial population responses. Surprisingly, the discriminability of stimuli near the adaptor increased, even though the responses of individual neurons to these stimuli decreased. This improvement arose mainly from how adaptation reshaped the geometry of the mean population responses, with changes in the noise structure playing a minor role. Locomotion modulated the population responses along a direction largely independent of adaptation, allowing the adaptation-induced increase in discriminability to persist during locomotion. Finally, artificial neural networks trained to reconstruct the visual stimulus under metabolic constraints reproduced the increase in discriminability and the decrease in responses. The paradoxical effects of adaptation are thus consistent with the efficient coding framework, allowing the brain to improve the representation of frequent stimuli while limiting the associated metabolic cost.

## Introduction

Visual perception is profoundly affected by adaptation to previous stimuli, which can induce changes in our ability to detect and discriminate stimuli ^1,2^, as well as generate visual illusions such as the tilt aftereffect ^3^. Adaptation-induced changes in perception have been connected to changes in visual responses ^4^ and have been observed in different species ^5–7^, sensory systems ^6,8^, and at different stages of sensory processing ^5,6,9,10^. Despite these studies providing a link between changes in perception and neural responses, the crucial question of how adaptation reshapes the geometry of neural representations remains largely unexplored. Understanding adaptation through this lens could reveal how adaptation reorganizes the distances among stimulus representations in population activity space—changes that determine discriminability and ultimately perceptual performance. Thus, what is still missing is a population-level, geometrical framework capable of relating adaptation-induced neural changes to their perceptual consequences.

A connection between adaptation-induced changes in perception and visual responses was obtained in studies of tuning curves in single neurons ^11^ or neural populations ^12^. These studies showed that adaptation can help neurons discriminate between stimuli ^11^ and decorrelate tuning curves ^12^. However, it is not clear whether all these observed changes are concerted to increase the ability of a downstream population to discriminate stimuli. Indeed, some of the single neuron response properties might be correlated at multiple levels: they could exhibit signal correlations (across different stimuli) and noise correlations (across multiple trials of the same stimulus) that can clearly affect the discrimination performance. In particular, any realistic readout of a population of neurons should generate a response for each stimulus every time it is presented (in a single trial). Noise correlations across different neurons might play an important role, and the average response properties of individual neurons, like their tuning curve, can only partially predict the discrimination performance. The tuning curve is typically estimated by looking at multiple trials, and hence, it is an average property of the neuronal response that cannot be read out in a single trial ^13,14^. Whether adaptation acts on the mean responses or on their variability or both is not obvious, and distinguishing them requires a single trial analysis of the discrimination ability. Finally, the typical tuning curves of individual neurons are affected in multiple ways by adaptation (e.g. by the stimulus presented in the present and by the activity of neurons driven in the past, see ^12^), and it is often difficult to capture all these changes using a simple model of the responses of individual neurons. Needless to say that all these multiple changes affect discriminability.

A more interpretable description of the activity of a neuronal population can be obtained by considering its representational geometry, i.e. the arrangement of all the points in the neural space corresponding to the different experimental conditions ^15^. This description of the data is interpretable in terms of perception and discriminability because it directly relates to the ability of a linear readout to discriminate between similar stimuli. Moreover, unlike with tuning curves, this description of the data can be applied to single trials. It might, therefore, reveal how adaptation-induced changes in representational geometry interact with other forms of modulation, such as those induced by running. Running strongly affects the visual responses ^16,17^ and their geometry ^18^, and reflects changes in behavioral state that might, in turn, affect adaptation ^19^. Overall, only a few studies have focused on adaptation-induced changes at the population level, and they often lacked trial-to-trial resolution ^20–22^, single-unit resolution ^12^. Among those that achieved both, analyses typically focused on changes in discriminability between pairs of stimuli, rather than examining how adaptation reshapes the full geometry of population responses across multiple stimuli ^7,23–27^. Applying a geometrical approach at the single trial level can reveal how adaptation depends on orientation similarity and whether adaptation-induced and running-induced changes interact in the neural code.

Furthermore, describing adaptation-induced changes from a geometrical perspective could shed light on the computational advantages of adaptation. Computational theories ^4,28^ have proposed that adaptation can efficiently represent stimuli through population homeostasis maintenance ^12^, optimization of information transmission ^29^, decorrelation ^30^, response-product homeostasis ^31^, and the trade-off between precision of the representation and metabolic cost^32,33^.

For all these reasons, we investigated how adaptation modifies the geometry of representations and how it affects the ability of a linear readout to discriminate between similar stimuli on single trials. We presented to awake mice sequences of visual stimuli in the form of oriented gratings characterized by two distributions: uniform and biased ^12^. As a first step towards answering which orientations become more discriminable, we first found that in the uniform environment, vertical orientations lead to the lowest discriminability, in contrast with cat ^34^ and primate ^35^ V1 neurophysiology, where visual acuity is lowest for oblique orientations. Then, we found that in the biased environment, discriminability increases for stimuli near the adaptor, consistent with human psychophysics studies ^2,36^ (but see ^37^), while responses near the adaptor decreased. This scenario is consistent with earlier phenomenological theories of adaptation ^38^. We found that this increase in discriminability arose mainly from a reshaping of the geometry of the mean responses and not so much on changes in the noise. We also observed that running expanded the geometry of representations. Running had a stronger effect in magnitude than adaptation on the geometry of neural representations. However, the increase in discriminability and decrease in responses around the adapted orientation was observed across different locomotion states.

Finally, we leveraged these data to constrain a theoretical model that predicts changes in discriminability in single trials, and that reveals a computational role for the response changes induced by adaptation. Following an efficient coding approach ^39,40^, we trained an artificial neural network to reconstruct stimuli presented in environments with the same distributions as those used in our experiments (uniform and biased). The network minimized a stimulus reconstruction cost and a metabolic cost to represent the stimuli efficiently. Consistent with our experimental data, we observed an increase in discriminability and a decrease in responses around the adaptor. Our model thus suggests that population responses to stimuli efficiently adapt to the environment statistics.

## Results

We presented sequences of static gratings interleaved by blank stimuli to awake head-fixed mice that were free to run on an air-suspended ball (Fig. 1a). The orientation of the gratings was sampled from a uniform distribution (Fig. 1b). We simultaneously recorded hundreds of neurons using two-photon imaging in layers 2/3 of V1 in these mice (Fig. 1c) ^16^.

**Figure 1.**
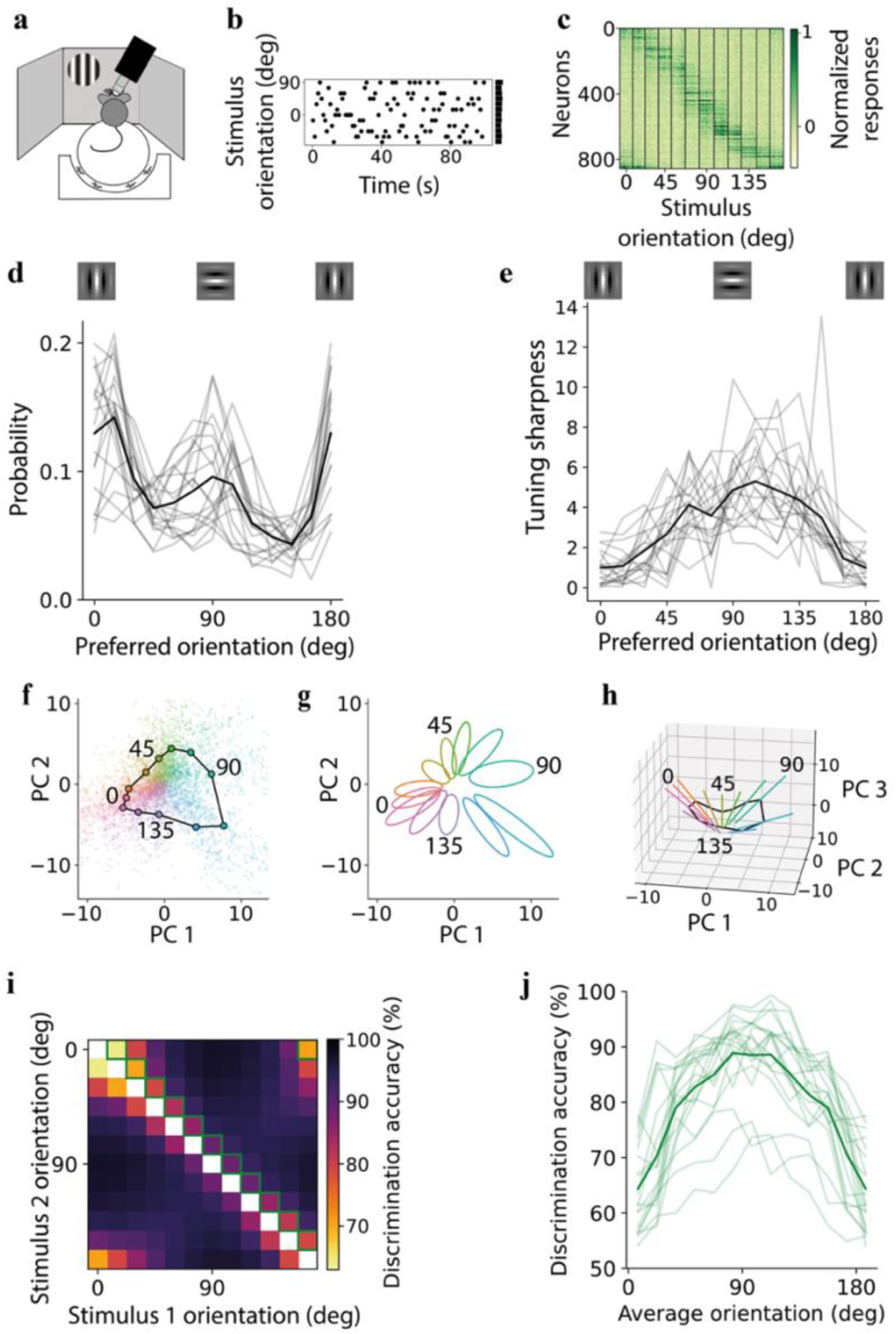
Heterogeneous single-neuron tuning shapes the geometry and discriminability of V1 population responses. **a**) Two-photon recordings of V1 neurons in head-fixed mice freely moving on a ball while viewing static gratings of different orientations. **b)** Example segment of oriented grating sequences drawn from a uniform distribution. **c**) Average normalized activity of all neurons recorded in an example session sorted by their preferred orientation. Orientation at 0 deg is vertical, while at 90 deg is horizontal. **d)** Distribution of preferred orientations across neurons for individual recording sessions (gray) and averaged across sessions (black). **e**) Orientation tuning sharpness (concentration parameter of von Mises functions fitted to the tuning curve of each individual neuron) as a function of preferred orientation, taken as the median across all neurons in each preferred-orientation bin, for individual recording sessions (gray) and averaged across sessions (black). **f**) The first two Principal Components (PCs) of the population responses in (c), showing responses in each repeat (small dots) and their averages for each stimulus (large circles). **g**) Covariance across trials of the population responses. For each stimulus orientation, the ellipse is centered on the average responses, and its axes are proportional to the square root of eigenvalues of the stimulus-conditioned covariance matrix of the trial responses in PC space. **h**) The first three PCs of the population responses, with lines indicating the main axis of the ellipsoid in (g) in three dimensions. **i**) Discrimination accuracy of any pair of stimuli, defined as discrimination accuracy of a linear classifier, averaged across all recording sessions. **j**) Discrimination accuracy for pairs of orientations differing by 15 deg (squares outlined in green in (i)) for each recording session (light green) and averaged across all sessions (dark green).

As a first characterization of how oriented gratings are encoded in the neural population, we studied the neurons’ tuning curves. We estimated the distribution of preferred orientations in the population of recorded neurons by calculating the circular mean of responses in an environment defined by a uniform distribution of orientations (Fig. 1d). Consistent with previous reports in mammals ^34,35,41^, we found an overrepresentation of neurons with a preference for the horizontal (90 deg) or vertical (0 deg) orientations (no significant difference in fraction of neurons with a preference for horizontal or vertical orientations across experiments: p = 0.09, Wilcoxon signed-ranked test). A larger number of neurons with a preference for 90 or 0 deg suggests a higher signal-to-noise ratio for these orientations, which appear consistent with a reduced discrimination threshold at horizontal and vertical orientations in human psychophysics experiments^1,34,42^. We also characterized the tuning width of the neurons by fitting von Mises functions to the responses of neurons to estimate the concentration parameters (high concentration corresponds to sharper tuning curves) (Fig. 1e). We found sharper tuning curves in neurons preferring horizontal orientations but wider tuning curves in neurons preferring vertical orientations. This dependence of tuning sharpness on angle was also present when considering only well-tuned cells or superneurons (Fig. S1d-f; one superneuron’s tuning curve is the average tuning curve of neurons that approximately prefer one specific orientation out of the 12 possible ones considered in this study). In conclusion, different tuning curves’ features (distribution of preferred orientations and tuning sharpness) suggest different patterns of orientation-dependent discriminability in the neural population.

To have a full picture, instead of considering single-tuning curves separately, we next focused on the full-dimensional neural activity space (i.e. the space in which the coordinate axes represent the activities of the different neurons). We projected the population responses into low-dimensional space and quantified distances in the full-dimensional activity space. The representations of the stimuli in the activity space reflected the circular symmetry of the visual stimuli (Fig. 1f), but as expected from the inhomogeneities in the tuning curve distribution and properties (Fig. 1d,e), they were not spaced uniformly around a circle. Instead, the distance in the activity space between stimuli with a horizontal orientation or similar was larger than for stimuli with a vertical orientation or similar. This can be seen in visualizations of the geometrical structure of the representations (Fig. 1f,g,h), which we created by using PCA to reduce the dimensionality of the activity space. The same results are also observed in the original full-dimensional activity space by comparing the Euclidean distance between stimuli near the horizontal orientation with the distance between those near the vertical orientation (p < 10^-5^, Wilcoxon signed-ranked test; Fig. S1a,b).

To characterize the geometry of the neural representations in a way that directly reflects the information that can be read out by a downstream population, we then measured the ability of linear decoders to discriminate pairs of stimuli (Fig. 1i,j), starting from the case in which all the orientations are presented with equal probability (uniform distribution). The performance of these decoders depends on the representations in the original full-dimensional activity space and, importantly, on the noise’s strength and structure. In general, we observed a simple relation between discriminability and Euclidean distances: the larger the distances, the more discriminable the stimuli (Fig. S1c). When considering pairs of stimuli 15 deg apart, we consistently observed a peak in discrimination accuracy around the horizontal orientation, while the lowest decoding performance was seen for the vertical orientations. These observations suggest that discriminability is primarily modulated by the distances between orientations, rather than by the size or the structure of the noise. Indeed, the noise level of horizontal stimuli was not clearly distinguished from the noise level of vertical stimuli (p = 0.21, Wilcoxon signed-ranked test; Fig. S1a,b). Moreover, the position of the discrimination peak did not depend on the noise structure, as pairwise correlations did not depend strongly on stimulus orientation (Fig. S2a) and removing correlations modestly increased the discrimination accuracy but not the peak position (Fig. S2b-d). Shorter Euclidean distances at vertical orientations are consistent with wider tuning curves at those orientations (Fig. 1e).

To understand the contribution not only of the tuning width but also of the distribution of preferred orientations to the orientation-dependent discriminability, we considered a lower-dimensional phenomenological model constructed from the average tuning curves of neurons in each of the 12 bins of preferred orientation. We fitted a von Mises function to each of these average tuning curves and then multiplied the fitted curves by the probability density of the associated preferred orientation (Fig. S3c). Alternatively, we only considered the preferred orientation distribution (Fig. S3a) or only the tuning width (Fig. S3b). We then considered the Euclidean distance between pairs of stimuli in the lower dimensional space of 12 superneurons with responses determined by these average tuning curves (Fig. S3d-f). When considering nearby stimuli (Fig. S3g-i), we could observe the decoding peak at the horizontal orientation only when taking into account the tuning width. However, the distribution of preferred orientations was also informative when considering the relation between discriminability and distances across all pairs of orientations (Fig. S3j-l). Hence, we concluded that both the tuning width and the response magnitudes contribute to the peak of discriminability at the horizontal orientation.

### Adaptation decreases responses of neurons tuned to the adaptor orientation while increasing discriminability around the adaptor

To understand how adaptation changes the individual neural responses and the geometry of neural representations, we presented sequences of oriented gratings with different statistics (Fig. 2a): stimulus orientation was sampled from either a uniform or a biased sequence ^12^, defining two environments that are characterized by different stimulus distributions. In the case of a biased sequence, one orientation was presented 50% of the time, which produced strong adaptation effects around that orientation. When projecting the activity in a space with reduced dimensionality, we observed that adaptation changes the representational geometry in a structured way (Fig. 2b). This geometry in the reduced activity space indicates that the discriminability might actually increase around the adaptor.

**Figure 2.**
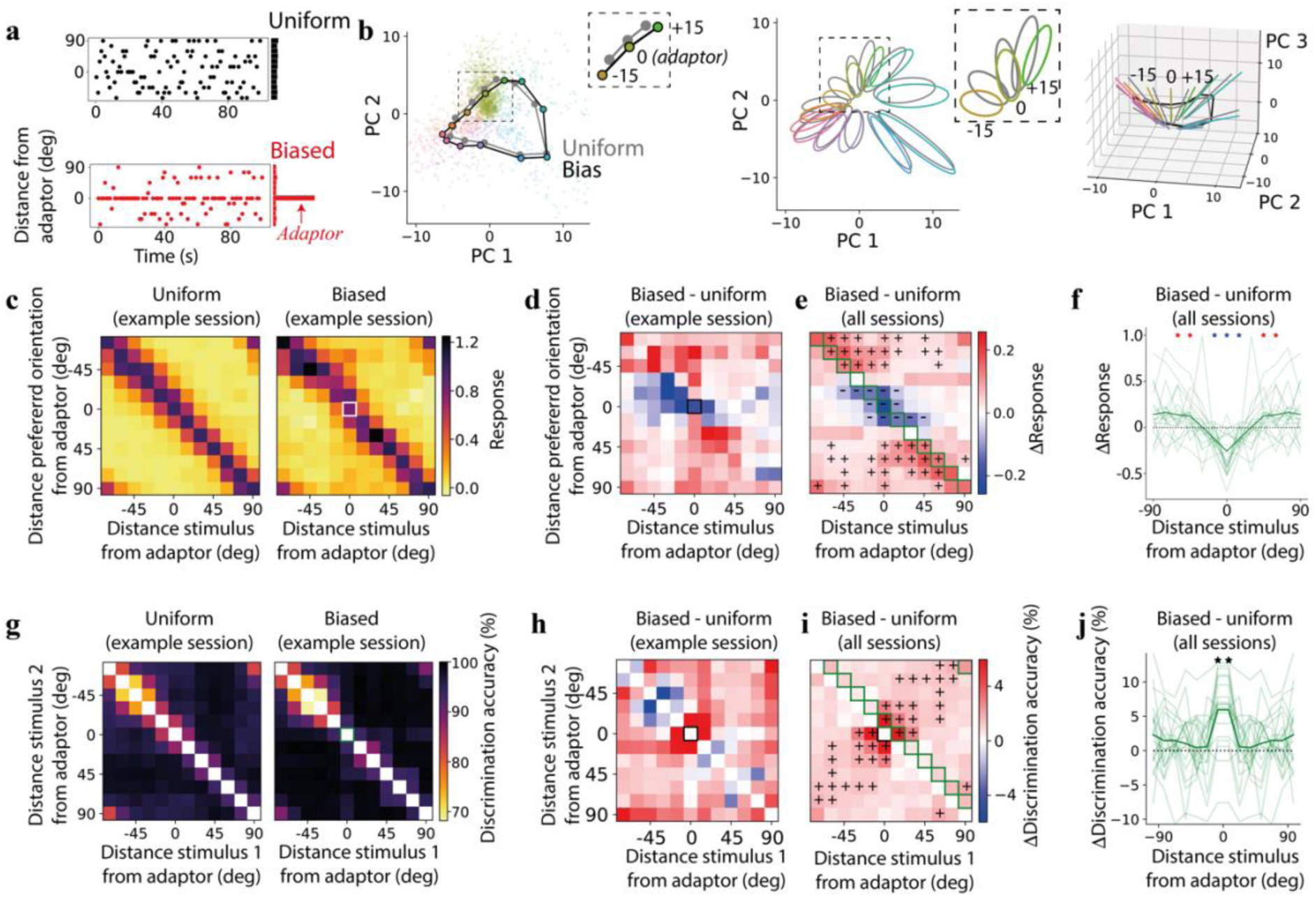
Adaptation increases discriminability around the adaptor while reducing neural responses. **a**) Example segment of oriented grating sequences drawn from a uniform (black, same as in Fig. 1b or biased distribution (red). **b**) Projection of neural activity in PC space, similar to Fig. 1f, but in a biased environment (colored dots, black lines), gray lines and dots correspond to the uniform environment; insets in the left and center panels focus on the adaptor orientations and orientations that are 15 deg distant from it. **c)** Average responses of tuned neurons in a single recording session (same as in panel b) whose preferred orientation has a given distance from the adaptor orientation (y-axis) to stimuli with a given distance from the adaptor orientation (x-axis); left: uniform environment; right: biased environment; white square: response to the adaptor stimulus of neurons tuned to the adapter. **d**) Difference in normalized responses between biased and uniform environments in (c). **e**) Same as in (d) but across recording sessions; pluses (resp. minuses) correspond to a significant increase (resp. decrease) in responses (p < 0.05, 1-sample t-test); green squares correspond to average values in (f); **f**) change in average responses between biased and uniform environments at the preferred orientation of each group of neurons (light green: single recording sessions; dark green: average across sessions; red asterisks: significant increase in responses as in (e); blue asterisks: significant decrease in responses as in (e)). **g**) Discrimination accuracy for any pair of stimuli in a uniform environment for one example session; right: same as the left panel but in a biased environment; green square corresponds to adapter orientation; **h**) Difference in population discrimination accuracy in an example recording between the biased and the uniform environment in (g). **i**) Similar to the example session in (h) but averaged across sessions; green squares correspond to average values in (j); pluses correspond to a significant increase in discrimination (p < 0.05, 1-sample t-test). **j**) Change in discrimination accuracy between biased and uniform environments for stimuli 15 deg apart as a function of the distance of the stimuli from the adapter; light green: individual recording sessions, dark green: average across all sessions; asterisks are a significant increase in discrimination accuracy (p < 10^-4^, 1-sample t-test).

We started by investigating how the tuning properties of neurons change in a biased environment. Consistent with previous reports ^12^, when averaging across recording sessions, we observed that adaptation induced a response decrease in neurons tuned to stimuli near the adaptor orientation (Fig. 2c-f, S4a,b, S7a). We also observed a response increase in neurons whose orientation preference was farther from the adaptor. In the original full-dimensional space, the Euclidean distance between the response to a given orientation and one 15 degrees away had the highest increase for the adaptor orientation (Fig. S4c,d).

We then compared the discrimination accuracy in a uniform environment to that in a biased environment. After estimating the accuracy in the two environments in the full-dimensional neural space (Fig. 2g), we computed the difference between the two environments (Fig. 2h,i, S7b). When considering pairs of similar stimuli that were 15 deg apart, we observed increased discrimination accuracy around the adaptor (Fig. 2j; p < 10^-4^, 1-sample t-test). This result was consistent across different adaptor orientations (adaptor at 0 deg: p < 10^-4^, 1-sample t-test; adaptor at 45 deg: p = 0.026, 1-sample t-test; Fig. S4e) as well as with increased cosine distance around the adaptor (Fig. S4f). Overall, changes in discriminability were qualitatively similar to observations in human psychophysics experiments ^2,36^ (but see also ^43^).

The increase in discriminability near the adaptor was driven by changes in mean responses rather than changes in variability. The standard deviation of single-neuron responses changed together with the tuning curves, decreasing for neurons tuned near the adaptor (Fig. S5a,b). To test whether this contributed to the changes in discriminability, we recomputed discrimination accuracy after rescaling the responses of each neuron to enforce a constant standard deviation, independent of stimulus and environment, while preserving the mean responses. The increase in discriminability around the adaptor persisted, with only a mild attenuation (Fig. S5c,d). We then measured pairwise noise correlations directly. Once the number of trials was balanced across conditions, their distributions were indistinguishable between the uniform and the biased environments (Fig. S5e), and their average changed significantly only for a few orientations far from the adaptor (Fig. S5f). The increase in discriminability around the adaptor was similarly robust when correlations were removed by shuffling the activity of each neuron independently across trials for a given stimulus condition (shuffling, see ^13,14,44^, Fig. S5g,h). The fact that the noise structure did not play a major role in adaptation-induced changes is consistent with previous reports ^12^ (but see: ^23^).

The increase in discriminability was already present in a low-dimensional description of the geometry. Because our intuition for the population geometry rests on projections onto the first two principal components (Fig. 2b), we asked whether such projections retain the changes measured in the full-dimensional space. Discrimination accuracy computed on an increasing number of principal components approached the accuracy obtained with all neurons at approximately eight components (Fig. S6a). Using only the first two components, we observed an increase in discriminability around the adaptor (Fig. S6b), which was also present with the first eight (Fig. S6c). Thus, discrimination accuracy keeps improving up to about eight components, showing that the population geometry is high-dimensional, while the first two principal components already convey the correct intuition that adaptation expands the representations around the adaptor (Fig. 2b), even though the underlying changes may extend beyond that plane.

In the previous analysis, the decoder was aware of changes in environments ^45^. We then asked whether a decoder unaware of changes in environments can deal with the changes in representational geometry ^45^. We trained linear decoders in a uniform environment and tested them in a biased environment or vice versa (Fig. S8a). The discrimination accuracy for pairs of approximately opposite stimuli relative to the adaptor orientation typically increased, including those near the adaptor (Fig. S8a). In other words, it would increase for pairs of stimuli whose orientation was approximately ±*θ* from that of the adaptor. Although the structure of the changes was not exactly the same compared to when a decoder was trained and tested in the same environment, there was a common result: near the adaptor, discriminability increased in biased environments, which was an even more surprising result when the decoder was unaware of a change in context. We also trained a regression model in a uniform environment to estimate the orientation of the stimulus presented (Fig. S8b). We confirmed a classical result showing, for stimuli near the adaptor, a repulsion of the estimated orientations away from the adaptor orientation ^6^ when tested in a biased environment.

Representations might change over time for reasons independent of the specific task we are considering (’representational drift’ ^46,47^). As the biased sequence followed the uniform one, one might wonder whether this geometric change could be explained simply by the temporal separation of the two environments. We thus asked whether we could discriminate the same orientation across two time blocks. The two time blocks could have the same statistics (uniform) or different statistics (uniform and biased). Discriminating between two blocks of time with uniform statistics was larger than chance (Fig. S8c,d). However, the discrimination accuracy was greater when discriminating between two blocks of time with different statistics. This difference in discrimination accuracy indicated that the passage of time alone was insufficient to explain changes in geometry (Fig. S8c,d).

We generalized the previous analysis by training and testing linear decoders in environments separated by different periods of time. Training and testing could be in the same (uniform) or different (uniform vs. biased) environments, but always separated by varying time blocks. As expected, given the potential presence of representational or recording drift, discrimination accuracy decreased over time (Fig. S8e).

Adaptation-induced changes in geometry emerged gradually within the biased environment and outlasted it. The two uniform blocks were separated by the biased one, and responses in the second uniform block retained the signature of adaptation present in the preceding biased block: relative to the first uniform block, neurons tuned near the adaptor responded less to stimuli near the adaptor, and neurons tuned far from it responded more (Fig. S9a), as in the biased environment itself (Fig. 2e). Discrimination accuracy (Fig. S9b) and cosine distance (Fig. S9c) near the adaptor also remained elevated in the second uniform block relative to the first, even though the environment was uniform. To resolve how these changes arose, we tracked cosine distances, which, unlike discrimination accuracy, can be estimated reliably from a few trials. Within the biased block, the cosine distance between the adaptor and the stimulus 15 deg away increased over approximately the first 8 minutes and then converged to a steady state (Fig. S9d), separating the representations near the adaptor by the end of the block (Fig. S9e).

### Locomotion and adaptation reshape the population geometry along largely independent dimensions

Running is one of the main drivers of visual response modulation in the mouse brain ^16,17,48^. Running increased the distance between stimuli (p < 10^-11^, Wilcoxon signed-ranked test; Fig. 3a). In the activity space, this expansion appears to be along directions that are approximately orthogonal to the dimensions spanned by all the visual stimuli (Fig. 3a), which is consistent with previous reports ^18,41^. This change in geometry enables the encoding of whether the animal is running or not without interfering with a linear readout of the orientation of the stimulus.

**Figure 3.**
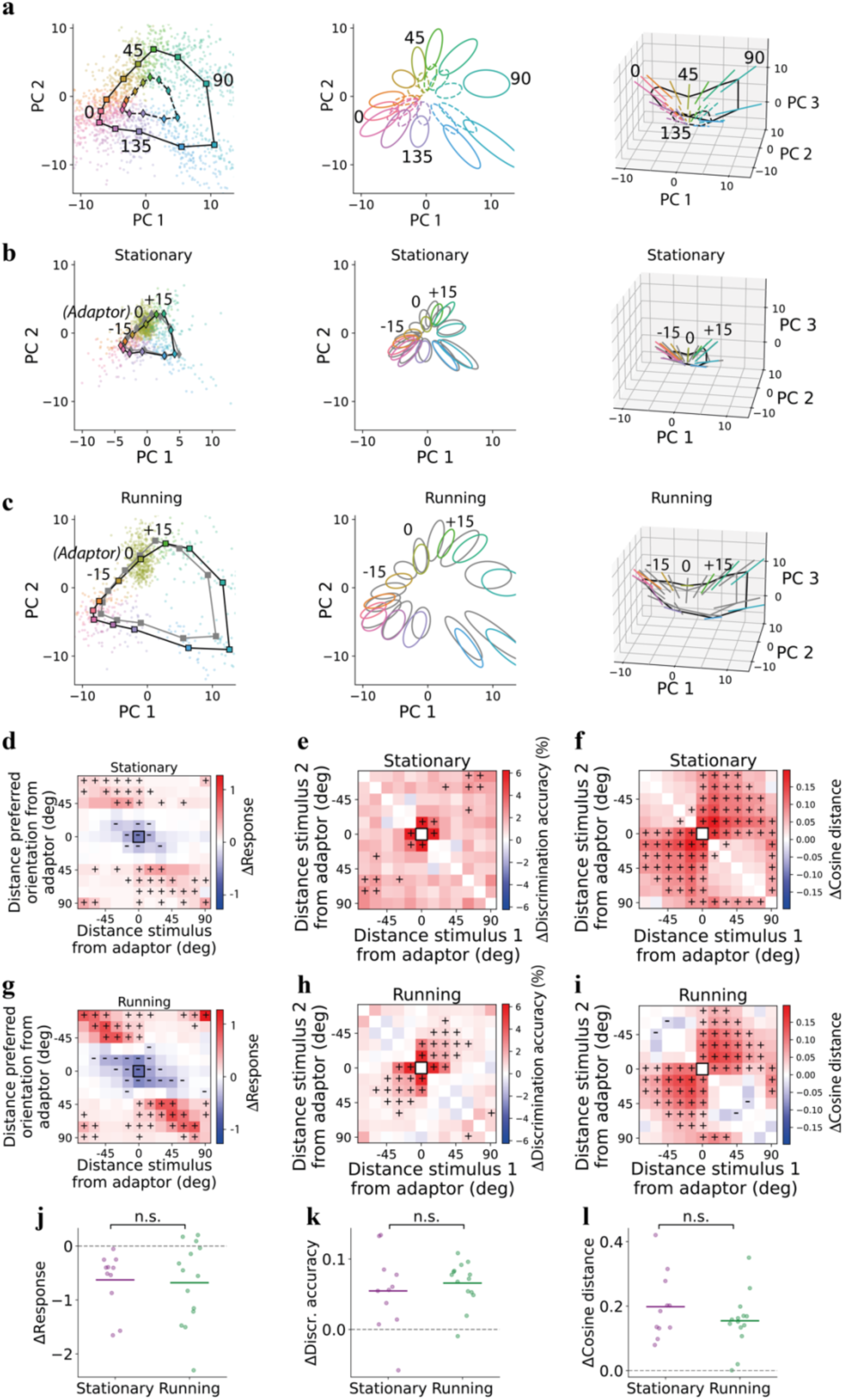
The adaptation-induced changes in geometry are preserved across stationary and running states. **a)** Projection of neural responses in a uniform environment in PC space similar to Fig. 1f-h, but comparing average stimulus responses measured when the mouse was running (squares, solid lines) vs. stationary (diamonds, dashed lines). Dots represent individual trials and are shown only during running. **b**) Projection of neural responses in PC space, similar to Fig. 2b, but for the stationary condition only. Colored dots and black lines correspond to a biased environment, while gray lines and dots correspond to a uniform environment. **c**) Similar to (b) but for the running condition. **d)** Difference in normalized responses between biased and uniform environments averaged across experiments similar to Fig. 2e, but only for the stationary condition. **e**) Difference in population discrimination accuracy between the biased and the uniform environment averaged across recording sessions similar to Fig. 2i but only for the stationary conditions; **f**) Change in cosine distance between angles from a uniform and a biased environment in the stationary condition; black square corresponds to adapter orientation; pluses correspond to a significant increase in discrimination (p < 0.05, 1-sample t-test); **g-i**) Similar to (d-f) but for the running condition. **j**) Change in normalized superneuron response at the adaptor orientation for stationary (black, n = 11 sessions) and running (red, n = 14 sessions) trials. Each dot represents one session. Horizontal bars indicate the mean. **k**) Change in pairwise discrimination accuracy for stimuli one step from the adaptor (0° and 15° from adaptor, symmetrized across ±15°). **l**) Change in cosine distance for stimuli one step diagonally from the adaptor (−15° and +15° from adaptor). In all panels, changes are computed as the adapted block minus the mean of the control blocks. Sessions are selected independently per locomotion condition (unbalanced). P-values are from two-sided Mann-Whitney U tests comparing stationary and running distributions.

Running strongly modulated neural activity, and the locomotion state accounted for more variability than the identity of the visual stimulus. We observed a general increase in Euclidean distances during running (Fig. S11a). We then computed the angle between the coding direction of running (i.e., the vector linking responses during the stationary condition to those during the running condition) and the main direction of variability (as measured by PCA) (Fig. S11b). We then compared this angle with the angle between the main direction of variability and the coding directions of different stimuli (the vector linking average responses to two orientations 15 deg apart, same locomotion condition). We found that the main direction of variability was more aligned with the running direction than the stimulus directions (Fig. S11c).

We compared the discrimination accuracy between any pair of stimuli for stationary and running conditions separately in a uniform environment. We observed a general increase in discrimination accuracy in the running condition (Fig. S10a), consistent with an increase in Euclidean distances (Fig. S11a). We then asked if the neural code for orientation is preserved across locomotion conditions. More specifically, we tested whether the coding directions for orientation (i.e. the vectors from one orientation to a different one) were parallel for running and stationary states. We computed the cross-condition generalization performance (CCGP) ^15,49^: we trained a linear model to discriminate any two angles in one locomotion condition (stationary or running) and tested it in the other condition (Fig. S10b). The CCGP (Fig. S10c) was similar to the performance achieved when we trained and tested the decoder using the same locomotion condition (Fig. S10a), supporting the notion that locomotion and stimulus representation in V1 are disentangled (or mostly disentangled ^18^).

We then asked whether the coding direction of running was preserved across stimulus orientations. We performed another CCGP analysis by estimating the ability to discriminate between stationary and running trials, training the decoder on one stimulus orientation, and testing on another orientation (Fig. S11d). We found it possible to decode locomotion, and we did not observe a large difference across angles. We also found that destroying correlations increased the discrimination accuracy of the previous analysis (Fig. S11e,f), which is consistent with the main axis of co-variability being aligned with running (Fig. S11b,c). These analyses indicate that there is a subspace in which the locomotion state of the animal is approximately invariant with respect to stimulus identity.

From the reduced-dimensional representations (Fig. 3a), it appears that the transformation of the geometry from stationary to running can be described as an expansion accompanied by a shift. It is then natural to ask whether this transformation can be explained by a simple scaling model in which all responses in the stationary case are simply multiplied by the same factor. We fit a scaling geometrical model (see Methods for more details), which can be described as a truncated cone, to the neural data. This model accounts only partially (averaged normalized error: 37%) for the change in responses by running (Fig. S11g). In conclusion, while our results on the scaling model indicate that the geometry is more complex than a truncated cone, those on CCGP show that expanding the geometry by running does not substantially interfere with the encoding of stimulus orientation.

Locomotion mostly preserved adaptation-induced changes. The analysis of the activity space with reduced dimensionality suggests that the changes in representations are approximately similar in different locomotion conditions (Fig. 3b,c). We then studied in which directions of the original full-dimensional neural space the locomotion-induced and adaptation-induced changes happen across orientations. For each orientation, we define the direction of adaptation as the vector between the points in the activity space that represent the responses in uniform and biased environments. We then estimated the angle between the coding direction of stationary/running and the direction of adaptation (Fig. S12a. This angle tended to be negative, although not significantly so. We also estimated the angle between the direction of maximum variability and the direction of adaptation (Fig. S12b). In this case, the angle was significantly negative, indicating that adaptation is oriented toward a decrease of this vector, which typically represents the average activity.

The coding directions for adaptation were preserved when the locomotion state changed. We computed the CCGP for adaptation by training the model to discriminate responses to the adaptor between the uniform and biased environment. We either trained the model in the stationary condition and tested it in the running condition or vice versa (Fig. S12c). We compared the CCGP with that obtained by discriminating between two uniform blocks, so that the only difference would be the passage of time. We found it easier to discriminate between biased and uniform environments than between two uniform environments (Fig. S12d). This shows that adaptation shifted the manifold in a direction not perfectly aligned with the locomotion axis. It would have been more difficult to discriminate between the two conditions if they had been perfectly aligned. When performing a similar operation—discriminating between stationary and running by training linear decoders in a uniform environment and testing them in a biased environment (Fig. S12e)—we did not observe a significant difference in the case of training and testing models in different uniform environments (Fig. S12f).

Finally, we tested how locomotion changes neural responses, discrimination accuracy, and cosine distances. We observed decreased responses (Fig. 3d,g; S12g,h) and increased discrimination accuracy (Fig. 3e,h) and cosine distances (Fig. 3f,i) around the adaptor during running and stationary periods. Across sessions, we did not find significant differences between the stationary and locomotion conditions around the adaptor (Fig. 3j-l). Thus, although running had a stronger effect on the geometry of representations than adaptation, we observed changes in both locomotion conditions that were similar to those reported without separating locomotion conditions (Fig. 2e,i).

### An artificial neural network reproduces the changes in discriminability and population tuning observed in mouse V1

Since theoretical work has shown that adaptation can increase the efficiency of neural representations, we next asked if our findings could be explained by efficient coding ^40^. We trained 6,000 autoencoder models with metabolic constraints (L1 norm in the hidden layer; Fig. 4a) and different hyperparameters (e.g. noise level, number of neurons, metabolic coefficient) to reconstruct the stimuli. We trained the autoencoder separately on uniform and biased statistics (Fig. 4b). In the biased environment, the autoencoder would be penalized more for misclassifying the adaptor’s orientation, as the adaptor is presented more often. The biased statistics also force the autoencoder to represent the adaptor in a more efficient manner than other orientations.

**Figure 4.**
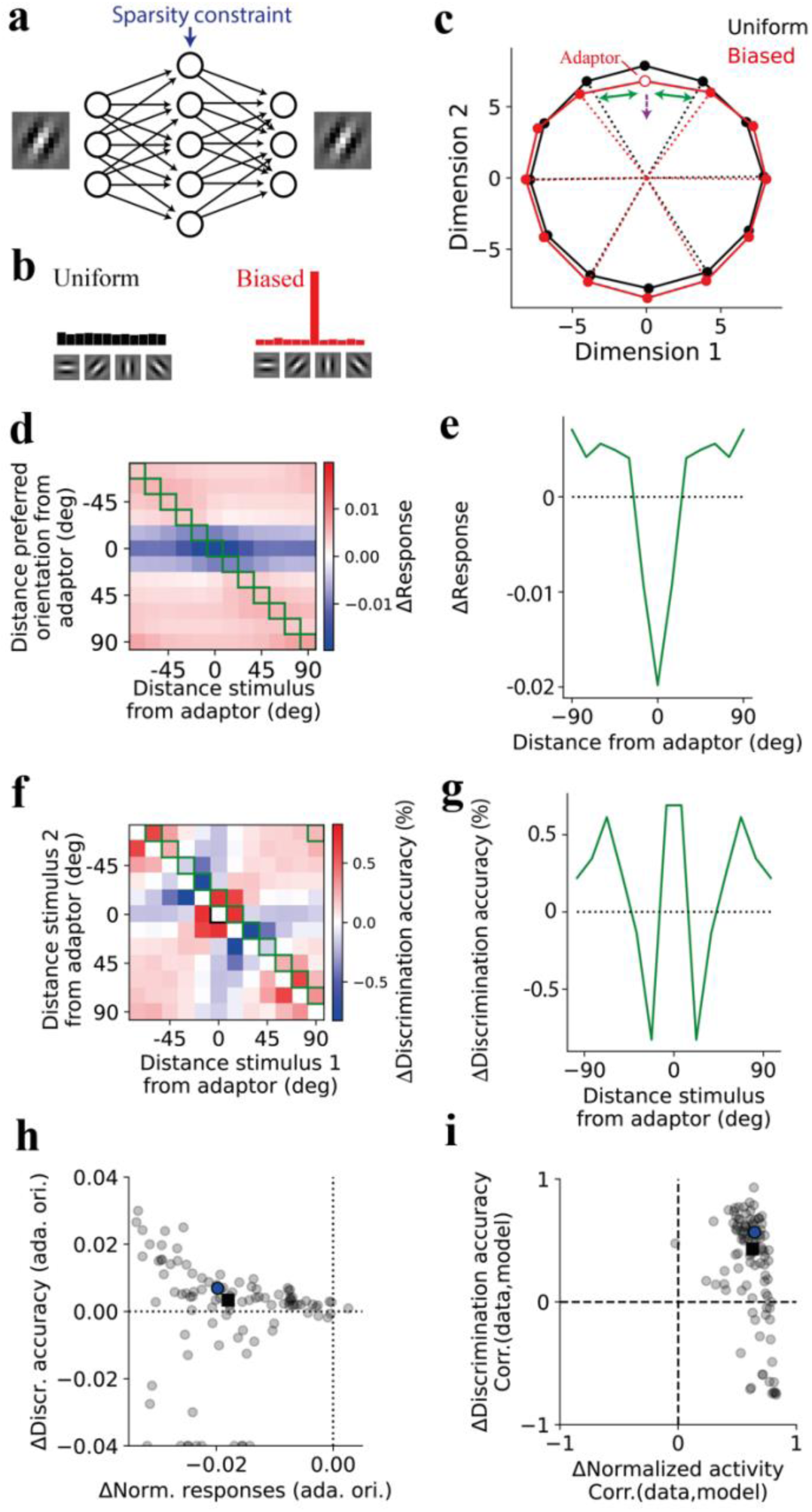
A normative model reproduces the changes in discriminability and tuning observed in mouse V1. **a)** Training set of the model matching the distribution of stimuli in the data. **b**) Autoencoder trained to minimize a multi-objective function composed of a reconstruction error in the output layer and energy cost (L1-norm sparsity) in the hidden layer; **c**) PCA on hidden-layer responses of one typical model, rotated with the adaptor on the vertical axis and the blank at the origin (compare with Fig. 2b and 5l). Purple arrow: response decrease near the adaptor; green arrows: distance increase. **e**) Change in average responses between biased and uniform environments at the preferred orientation of each group of neurons, computed as in Fig. S4b but for the example model in (c). **f**) Difference in discrimination accuracy in an example recording between the biased and the uniform environment, computed similarly to Fig. 2h but for the example model in (c). **g**) Change in discrimination accuracy between biased and uniform environments for stimuli 15 deg apart as a function of the distance of the stimuli from the adapter, computed similarly to Fig. 2j but for the example model in (c). **h**) Changes in discrimination accuracy near the adaptor as a function of changes in responses for neurons tuned to the adaptor at the adaptor orientation for different model’s hyperparameters. **i**) Correlations between data and models for the difference in discrimination accuracy and the difference in normalized activity between a uniform and a biased environment. Each circle corresponds to the average result across autoencoder models (n=30) with the same parameters. The blue circle is an example solution in (c-g), while the black square represents the median across all the models.

We assumed that inhomogeneities in tuning curves induced by adaptation can be explained separately from those in the “base” uniform conditions. We thus developed a network model whose symmetries would ensure that in the uniform conditions, tuning curves would be homogeneous, abstracting away from the inhomogeneities in the data. Despite the simplicity of the model and the minimal number of assumptions, adaptation changed the population geometry in most models in a way that is similar to the data (Fig. 4c). Within a broad hyperparameter region, we observed a decrease in responses around the adaptor for neurons tuned to the adaptor (Fig. 4d,e) and an increase in discrimination accuracy around the adaptor (Fig. 4f,g) consistent with our experimental observations (Fig. 2e,f,i,j).

Among the different hyperparameters of the model, we focused on the metabolic penalty and its impact on the metabolic cost, discrimination accuracy, and response magnitude (Fig. S13a-f). We observed that a decreased metabolic penalty increased response magnitudes (Fig. S13b) and discriminability (Fig. S13c) when averaged across all orientations or pairs of orientations. Furthermore, a decrease in metabolic penalty leads to a weaker (and negative) change in response magnitude (Fig. S13e) and an increase in change in discrimination accuracy (Fig. S13f) around the adaptor. These results can be visualized in reduced dimensions as an expansion of the geometry (Fig. S13g) and suggest that the running-induced changes could be interpreted as a decrease in metabolic penalty.

The decrease in responses and the increase in discriminability around the adaptor were robust across a wide range of hyperparameters. For each model, we compared the change in response at the adaptor with the change in discrimination accuracy near the adaptor. In most models, the two changes co-occurred, with responses falling and accuracy rising, as we observed in V1 (Fig. 4h, S13d). To compare model and data quantitatively, we correlated the model’s difference in discrimination accuracy and difference in normalized activity between the uniform and the biased environment with those measured in V1. Most models correlated positively with the data for both quantities (Fig. 4i). The example solution of Fig. 4c-g is therefore representative of the model class rather than a singular case.

We included blank stimuli in the training set to mimic the zero-contrast gray screen that interleaved the gratings in the experiments, but they were neither the source of the adaptation effects nor of the model’s sparsity. A model trained without any blanks still reorganized its geometry (Fig. S14a), decreased its responses near the adaptor (Fig. S14b), and increased its discrimination accuracy near the adaptor (Fig. S14c). Sparsity, quantified by the Gini index across neurons, increased with the fraction of blanks, but only modestly: it was already present when blanks were entirely absent (Fig. S14d).

The number of hidden units determined whether the model reproduced the population geometry. With too few hidden units, the projection of the responses was distorted (Fig. S15a), and adding units attenuated the distortion (Fig. S15b,c). Most units retained non-zero activation despite the L1 penalty, which may stem from the sigmoid activation function or from input noise: the Gini index increased with metabolic weight (Fig. S15d) and decreased with input noise (Fig. S15e). We read out 50 units because larger numbers saturated discrimination accuracy near 100% in both environments, masking the effect of adaptation (Fig. S15f).

The effects also did not depend on the fully connected architecture. Because nearby pixels in natural images are correlated, we trained convolutional autoencoders on the same task. A convolutional autoencoder using 1.7% of the parameters of the model in Fig. 4, reproduced the reorganization of the geometry (Fig. S16a), the decrease in responses near the adaptor (Fig. S16b), and the increase in discrimination accuracy near the adaptor (Fig. S16c).

### Reduced responses and enhanced discriminability near the adaptor reflect an efficient coding strategy

The decrease of responses by adaptation in neurons tuned to the adaptor orientation in the data (Fig. 2e) and the normative model (Fig. 4d) may appear at odds with increased discriminability around the adaptor observed in our data (Fig. 2i) and in the normative model (Fig. 4f). We now show that what we observed in the data reflects an interesting computational strategy that allows the system to better discriminate more frequent stimuli without increasing the metabolic cost. We will compare this strategy with two other scenarios: one where the adaptor discriminability increases but also the metabolic cost, and another where the energy consumption is reduced but also the discriminability of the adaptor decreases.

We started by considering a hypothetical adaptation of tuning curves where the peak responses of the neurons tuned to the adaptor were either depressed (Fig. 5a, S17a) or facilitated (Fig. 5b, S17b). We also considered another hypothetical adaptation of tuning curves (Fig. 5c) consistent with changes observed in the data (Fig 2h) but applied to homogeneous tuning curves. In this way, similarly to the computational model (Fig. 4), we would abstract our results away from the inhomogeneities we observed in a uniform environment (Fig. 1) and focus on the changes in a few response properties.

**Figure 5.**
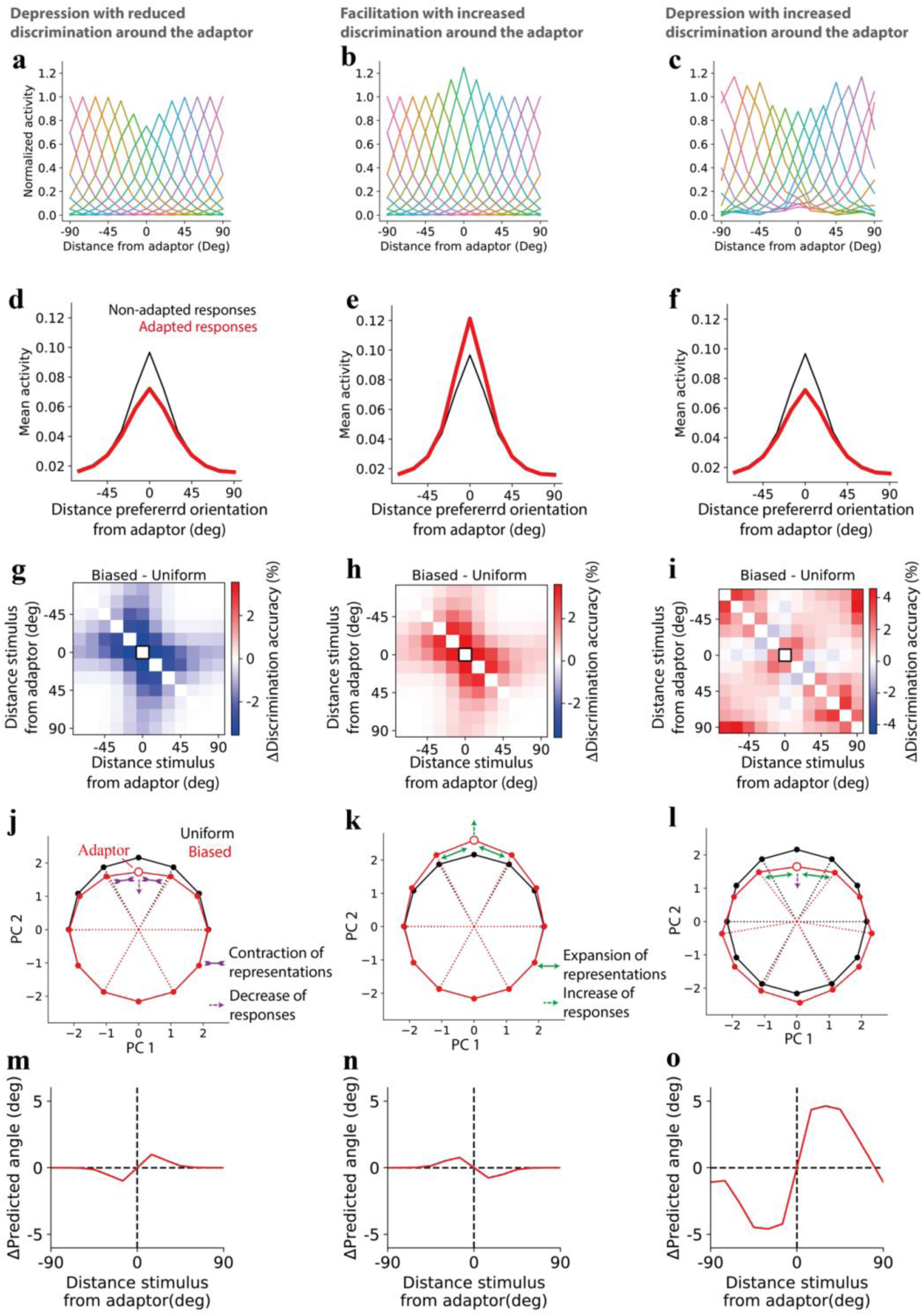
The empirically observed response change reshapes the geometry to reconcile reduced responses with increased discriminability, unlike a simple depression or facilitation. **a**) Normalized population-average responses to different orientations. Each curve corresponds to the average responses of neurons tuned to a specific orientation in a hypothetical, biased environment where the adaptor decreases responses in neurons tuned to the adaptor at the adaptor location. **b**) Similar to (a) but the adaptor increases responses in neurons tuned to the adaptor at the adaptor location. **c**) Similar to (a) and (b) but the increase and decrease in responses reflects that observed in the data (see Methods). **d-f**) hypothetical average firing rate of neurons if their responses were not adapted but the distribution of orientation was biased (black) and firing rate of neurons when their responses are adapted as in (a-c) consistently with a biased distribution of orientations (red). **g-i**) difference in discrimination accuracy (here, computed based on population discriminability between any pair of stimuli) between a homogenous population (representing biased condition) and the same population after applying the perturbation in tuning curves in (a-c). Black squares correspond to adapter orientation; **j**) Projected neural activity in PCA space before and after changes in (a) decreases responses (purple dashed arrow toward the center) and reduces distances and discrimination accuracy near the adaptor (purple inward arrows). **k**) Same as in (j), but changes in (b) are applied, which increase responses (green dashed arrow farther from the center) and enhance distances between stimuli near the adaptor (green outward arrows) as well as discrimination accuracy. **l**) Despite the response decrease (purple dashed line), the changes in (c) enhance distances and discrimination accuracy near the adaptor (green outward arrows). **m-o**) Angle prediction in the biased environments of (a-c) after training a model in a uniform environment. The angle is calculated from a linear regression of cos *θ* and sin *θ*, where *θ* is the stimulus orientation, followed by computing arctan(sin *θ*⁄cos *θ*).

What is the advantage of a simple response depression (Fig. 5a)? We measured the metabolic cost as the average firing rate over time based on the biased environment in the experiments. Assuming this biased environment, we compared the metabolic cost if the responses were uniform (non-adapted) or depressed (Fig. 5d). As expected, a response depression around the adaptor led to a decrease in metabolic cost compared to the non-adapted responses. We then estimated discrimination accuracy to different pairs of stimuli, assuming independent and identically distributed noise when responses were depressed compared to when were non-adapted. As expected, because of a decrease in signal-to-noise ratio, discrimination accuracy decreased around the adaptor, thus to the most frequent stimuli. Thus, response depression has the advantage of decreasing metabolic cost (Fig. 5d) but the disadvantage of decreasing discriminability (Fig. 5g). The reverse emerges when considering response facilitation (Fig. 5b): facilitation has the advantage of increasing discrimination accuracy around the adaptor (Fig. 5h) but the disadvantage of increasing metabolic cost (Fig. 5e).

Finally, we estimated the metabolic cost and discrimination accuracy as the average firing rate based on the biased distribution used in the experiments. What are the consequences of these tuning curve changes in a biased environment? The changes in responses observed in the data and applied to a homogenous population present two advantages: they decrease the metabolic cost (Fig. 5f) and, at the same time, increase discrimination accuracy around the adaptor (Fig. 5i).

To find an intuitive explanation for how these two seemingly incompatible changes can happen simultaneously, we inspected the representations projected in a reduced dimensionality activity space (Fig. 5l). We observed that while responses near the adaptor got closer to the center, as expected from a decrease in responses, the stimuli near the adaptor got farther away from it, leading to a local increase in discrimination accuracy. These results can be compatible with a combination of changes in gain and warping of tuning curves ^50^ and they can be contrasted to the cases of simple depression (Fig. 5j) or facilitation (Fig. 5k), in which there was an increase or a decrease in both responses and discrimination accuracy.

The adaptation-induced changes observed in the data and reproduced by the model have two benefits: reduction in metabolic cost and increase in overall discriminability. Is there any cost? Let us assume that the visual system decodes the stimulus orientation by estimating the angle from the neural population vector. Let us also assume that the decoder is unaware of changes in representational geometry ^45^ (Fig. S8a,b). This means that a decoder trained in a default environment (e.g. a uniform environment) could be tested in another environment (e.g. one specific biased environment) differently from a decoder trained and tested in the same environment (Fig. 2). Then, for the unaware decoder, there is a strong repulsion of the estimated angle near the adapted orientation (Fig. 5o), i.e. the estimated angle is farther from the adaptor than it really is (as it can be seen in the PCA space, Fig. 5l), creating a bias. This repulsion is what we observed in the data (Fig. S8b) and is consistent with a stronger reduction of tuning curves at the flank near the adaptor ^12^.

Repulsion of orientations near adaptation is consistent with the tilt aftereffect ^38,51,52^, where a long exposure to a stimulus (e.g., vertical) can produce the illusion that a different but similar stimulus (e.g., slightly oblique) will be perceived more different than what it is (e.g., more oblique). This issue would be less prominent in the other hypothetical adaptation-induced changes in geometry (Fig. 5j,k,m,n), suggesting that having an unbiased representation of the orientation may be, to some degree, less important than a robust discriminability between two angles and a metabolic cost under control. In other words, adaptation-induced changes may improve the ability to tell apart two stimuli (focusing on their relative difference of orientations) while decreasing the ability to identify the absolute orientation of those stimuli.

## Discussion

We investigated how the representational geometry in mouse V1 is shaped by adaptation under environments with different stimulus statistics. More specifically, we presented oriented gratings sampled from uniform and biased distributions. Consistently with previous studies in cat V1 ^12^, we observed that adaptation decreases the neurons’ responses to the adaptor orientation. However, we also observed increased discriminability around the adaptor. A normative model could explain these results, suggesting that the observed changes in the representational geometry emerge from a trade-off between improving the representation of frequent stimuli and reducing the metabolic cost of responding to these more frequent stimuli.

Adaptation-induced changes in discriminability have been reported across multiple brain areas, including V1 ^7,24,27^, V2 ^27^, V4 ^25^, and MT ^26^, as well as across different species such as macaques ^25–27^, marmosets ^24^, and mice ^7^. However, none of these studies examined changes in the statistics of the environment and, in general, changes in the geometry of neural representations, as we did in our study.

The analysis of the geometry in the uniform case is not fully consistent with human psychophysics studies. In humans, visual acuity is higher for both horizontal and vertical orientations and lower for oblique stimuli ^1,53^, a phenomenon known as “oblique effect” ^42^, which is consistent with several neurophysiology studies in cat ^34^ and primate V1 ^35^. However, we found a bias only towards horizontal orientations, while vertical orientations were less discriminable than both horizontal and oblique orientations. The reason for this discrepancy is not clear. One possible explanation is that, in the natural scene observed by mice, especially while running, vertical shapes are less frequent or less important, a prediction that can be tested experimentally.

The geometric analysis in the biased case revealed that, despite the decrease in responses, discriminability increases between the stimuli around the adaptor, consistent with human psychophysics studies ^2,36^ (but see ^37^) and theoretical models ^36,54^. A reduced dimensionality analysis shows intuitively how decreases in responses and increase in discriminability can coexist through a non-uniform transformation of the circle representing the uniform environment. The geometry in the low-dimensional space is consistent with earlier phenomenological theories of adaptation ^38^ but was not until now shown in neural population data. In previous studies in primate V1 ^11^ only an analysis at the level of single neurons was performed, except for a study on neuron pairs ^23^, which focused only on local discrimination and thus did not on the full population geometry. Furthermore, the results on pairwise noise correlations in primate V1 in ^23^ were different from those in cat V1 ^12^. It remains to be understood whether our population analysis of mouse V1 would match that of primates and cats.

Adaptation reshaped discriminability through the signal rather than the noise. A population can in principle discriminate better in three ways: by separating the mean responses^11^, by reducing the variability of individual neurons, or by changing the structure of correlations between neurons^23^. Our data point to the first. Single-neuron variability changed together with the tuning curves, yet enforcing a constant variability across stimuli and environments left the increase around the adaptor in place, with only mild attenuation (Fig. S5a-d); pairwise correlations differed little between environments, and significantly so only for a few orientations far from the adaptor (Fig. S5e,f). The direction of the noise points the same way: the principal axis of population variability is more aligned with the locomotion axis than with the stimulus axes (Fig. S11b,c), and the adaptation vector is anti-aligned with that axis (Fig. S12b), so adaptation reduces activity along the dominant direction of variability instead of reshaping it. Note that changes in the noise structure were reported to improve discriminability in macaque V1^23^, after adaptation lasting hundreds of milliseconds, leaving open whether the species or the timescale of adaptation determines which mechanism dominates. Each of these three mechanisms was a priori plausible. Our results attribute the change predominantly to the separation of the mean responses, restricting candidate mechanisms to those that modify tuning curves rather than the structure of response variability.

Furthermore, in an experimental setup in which stimulus latency was reduced to enhance adaptation effects, adaptation-induced improvements in discriminability in mouse V1 did not translate into clear improvements in task performance: the discrimination threshold increased, whereas false-alarm rates decreased with shorter latency. Given the difference in the experimental setup in ^7^ from our setup, where instead of reducing stimulus latency we modified probability distributions, it remains to be determined how changes in discriminability in the V1 population reflect changes in discrimination performance in behavioral tasks.

The peak in discriminability followed the statistics we imposed rather than those of the natural environment. In principle, the increase around the adaptor could have reflected a coincidental match between our biased distribution and the orientations that mice encounter most often, and so a pre-existing bias rather than a response to our manipulation. Our design excludes this possibility: we used two biased environments, with the adaptor at 0 or at 45 deg, and in each of them discriminability increased around the adaptor of that experiment (Fig. S4e). A fixed bias in the natural environment cannot produce a peak that moves with the adaptor.

Note that all comparisons were made between a biased environment and a uniform environment. This comparison shows an increase in responses for neurons whose preferred orientation is far from the adaptor (offset neurons). How would discriminability change if responses in a biased environment were instead compared to responses to an unadapted environment, where grating stimuli are preceded by prolonged blank stimuli? Such modulation could result in a lack of facilitation in the offset neurons. Consequently, this lack of facilitation would not improve discriminability. We believe this is consistent with our efficient coding framework. In an unadapted environment where gratings are sparse, as they are often preceded by blank stimuli, energy expenditure would be low most of the time, since responses to blank stimuli are low. Thus, the brain could afford strong responses to rare gratings, thereby improving their discriminability, which would be hard to further improve even around the adaptor in a biased environment.

Running strongly affects visual responses ^16,17,48^, and adaptation effects can depend on the animal’s behavioral state ^19^. We observed that, while running expands the stimulus representations, the increase in the discriminability of an adapted stimulus and the decrease in its responses are present during both stationary and running periods.

To understand how orientation, adaptation, and running information are formatted in V1, we performed several CCGP analyses ^15,49,55^. We found that the discriminability of a pair of stimuli depended on the test locomotion condition (higher during running). However, the coding directions for pairs of stimuli in the two locomotion conditions were approximately the same, enabling a decoder trained on stationary stimuli to work also in the running condition (and vice versa). The same geometry supported a higher-than-chance discriminability of stationary vs. running conditions when training a decoder in one orientation and testing it in another one, suggesting that the coding direction of running was approximately preserved across stimulus orientations. Another CCGP analysis suggested that it was also preserved across environments, and thus, adaptation did not affect the running direction. Finally, a different CCGP analysis showed that adaptation shifts the manifold in a direction not perfectly aligned with the locomotion axis.

All these results indicate that the locomotion state is encoded, and despite that, it is not strongly interfering with the readout of the stimulus orientation. In other words, stimulus orientation and locomotion states are two approximately disentangled variables, enabling a simple linear readout to generalize across multiple situations without any need for retraining. This indicates that the geometry is dominated by a relatively low-dimensional structure ^18,56^, which is not trivial to observe in activity spaces whose ambient dimensionality is elevated (i.e. when considering a large number of neurons). This structure has been observed in multiple brain areas across different species ^15,49,55,57,58^.

Normative theories of adaptation have been based on different frameworks, not necessarily mutually exclusive, including redundancy reduction, predictive coding, surprise salience, inference, and efficient coding ^28^. Following an efficient coding approach ^40,59^, we trained an artificial neural network ^39^ to represent stimuli in different environments under energy constraints. Variations of an efficient coding approach have considered different objective functions, such as maximizing mutual information ^60^. Here, similarly to previous work, we considered a tradeoff between representation fidelity and metabolic cost ^32,39,61^. Differently from previous studies, we not only considered its effect on changes in tuning curves and perceptual effects but also on the full population geometry. After comparing networks trained under biased or uniform statistics, we observed an increase in discriminability and a decrease in responses around the adaptor, consistently with our experimental data. It would be interesting to understand how our theory would apply to different forms of adaptation, such as contrast adaptation ^20,22^ or even affecting other sensory modalities, such as the auditory one ^62^, and compare it to other related normative theories ^63,64^.

Several properties make the autoencoder well suited to early sensory areas. Its representations follow from the structure of sensory experience alone, which fits our experiment: the animals performed no task, leaving the input statistics as the main quantity available to shape the geometry. It also admits a local implementation^65^, placing the principle within reach of a biological circuit, though adaptation in V1 likely also involves interactions with other areas. In both respects, the model stays agnostic about implementation and about time, as in a long tradition of efficient-coding models ^59,60,66,67^. Adaptation itself unfolds over multiple timescales, and the changes we report build up gradually within the biased environment and persist beyond it (Fig. S9); models that track environmental statistics continuously could capture this time course. What the autoencoder model does show is that discriminability around the adaptor improves in a network whose loss rewards only reconstruction and energy constraints — better discrimination of frequent stimuli emerges from efficient coding alone, without ever being an objective.

In conclusion, our model suggests that the stimuli representation is efficiently encoded in a way that considers the stimulus statistics. Several open questions stem from our study. The first question is to understand the detailed neural mechanisms underlying the observed phenomena in the data. The second question is whether the mouse perception reflects the finding in the neural population. Answering this question will require the animal to perform a discrimination task.

## Acknowledgments

This work was supported by NIH (R21 EY035064 to MD and DLR and U19 NS107613 to KDM); NSF (NeuroNex Award 1707398 DBI to KDM, and SF); the Gatsby Charitable Foundation (GAT3708 to KDM, and SF); the Simons Foundation to SF; the Swartz Foundation to SF and KDM; Kavli Foundation to MD, SF and KDM; and the Wellcome Trust (223144/Z/21/Z to MC and KDH). We thank Lauren Wool for her support in designing a set of visual stimuli used in this work. We also thank Joram Keijser and Xuexin Wei for helpful comments on the manuscript.

## Author contributions

MD and MC designed the experiments; MD and SB performed the experiments; SB and CBR performed surgery; MD, RN, and YF preprocessed the data; MD, RN, DLR, KDM, MC, and SF designed the data analysis; MD performed the data analysis; MD, RN, and SF designed the model; MD and RN developed the model; MD, RN, KDH, DLR, KDM, MC, and SF interpreted results; MD wrote the original draft; MD, RN, SB, DLR, KDM, MC, and SF edited the manuscript.

## Methods

All experimental procedures were conducted in accordance with the UK Animals (Scientific Procedures Act) 1986. Experiments were performed at University College London under personal and project licenses released by the Home Office following appropriate ethics review.

### Mice

We recorded neural activity from 12 transgenic animals (5 males, 7 females) in which specific cell types were labeled by a functional or structural indicator. In this study, we focused on all neurons recorded, independently of cell type. Experiments in which an interneuron class was labeled with tdTomato and recorded together with other cells were conducted in double-transgenic mice obtained by crossing Gt(ROSA)26Sor < tm14(CAG-tdTomato)Hze > reporters with appropriate drivers: *Pvalb*<tm1(cre)Arbr > (1 male, 1 female), *Vip*<tm1(cre)Zjh > (1 female), *Sst*<tm2.1(cre)Zjh > (3 males, 1 female), and GAD-nls-mCherry (1 male, 2 females). Experiments in which indicator was expressed uniquely in one neuron class were conducted in single transgenic mice: *Scnn1a-Cre* (1 female). Mice were used for experiments at adult postnatal ages (P59-214).

### Animal preparation and virus injection

The surgeries were performed in adult mice in a stereotaxic frame and under isoflurane anesthesia (5% for induction, 0.5%–3% during the surgery). During the surgery, we implanted a head-plate for later head fixation, made a craniotomy with a cranial window implant for optical access, and, on relevant experiments, performed virus injections, all during the same surgical procedure. In experiments where an interneuron class was recorded with other cells, mice were injected with an unconditional GCaMP6m virus, AAV1.Syn.GCaMP6m.WPRE.SV40 (#100841; concentration 2.23 10^12^). In experiments where a cell type (excitatory L4 neurons) was labeled by unique expression, mice were injected with AAV-Flex-hSyn-GCaMP6m (#100845; concentration 2.23 10^12^). In a subset of mice crossed with GAD-nls-mCherry (n = 2 females), a sparse set of unspecified neurons (most of them excitatory) were labeled, and the following viruses were injected: pAAV-FLEX-tdTomato (#28306-AAV1; concentration 2.5 10^12^); pENN.AAV.CamKII 0.4.Cre.SV40 (#105558; concentration 4 10^8^). All viruses were acquired from University of Pennsylvania Viral Vector Core. Viruses were injected with a beveled micropipette using a Nanoject II injector (Drummond Scientific Company, Broomall, PA 1) attached to a stereotaxic micromanipulator. Six to seven boli of 100-200 nL virus were slowly (23 nl/min) injected unilaterally into monocular V1, 2.1-3.3 mm laterally and 3.5-4.0mm posteriorly from Bregma and at a depth of L2/3 (200-400 mm).

### Visual stimuli

Stimuli were horizontal static two-dimensional Gabor functions presented in a location adjusted to match the center of GCaMP expression on one of two screens that spanned 45 to +135 of the horizontal visual field and ± 42.5 of the vertical visual field. During the gray screen presentation (duration 0.5 s), the screens were set to a steady gray level equal to the background of all the stimuli presented for visual response protocols. Gabor functions were presented for 0.5 s, with a spatial frequency of 0.1 cycles/deg and a width of 13 Deg.

### Stimulus environments

The static gratings presented in the sequences were sampled from either uniform or biased distributions. In the biased distribution, one orientation (45 deg, n=12 recordings; 0 deg, n=7 recordings) was presented 50% of the time. The other stimuli were sampled from a uniform distribution in the remaining trials. In n=6 recordings, the contrast of 7% of the repeats was set to 0, otherwise the contrast was always at 100% at the center of the Gabor stimuli. The number of trials in each sequence was approximately either 1,000 (n=1 recording), 2,000 (n=12 recordings) or 2,500 (n=6 recordings). We always presented first a uniform environment, then a biased environment and lastly another uniform environment.

### Imaging

Experiments were performed at least two weeks after the virus injection. We used a commercial two-photon microscope with a resonant-galvo scanhead (B-scope, ThorLabs, Ely UK) controlled by ScanImage ^68^, with an acquisition frame rate of about 30Hz (at 512 by 512 pixels, corresponding to a rate of 4.28-7.5 Hz per plane), which was later interpolated to a frequency of 20 Hz, common to all planes. Recordings were performed in the area where expression was strongest. In most recordings (n = 16) this location was in the monocular zone (MZ, horizontal visual field preference > 30 deg) ^69^. Other recordings (n = 11) were performed in the callosal binocular zone (CBZ, n = 4, 0-15 deg) ^70^ and others (n = 7) in the acallosal binocular zone (ABZ, 15-30 deg).

### Data analysis

#### Data processing

We analyzed raw calcium movies using Suite2p, which performs several processing stages ^71^. First, Suite2p registers the movies to account for brain motion, then clusters neighboring pixels with similar time courses into regions of interest (ROIs). Based on their morphology, we manually curated ROIs in the Suite2p GUI to distinguish somata from dendritic processes. For spike deconvolution from the calcium traces, we used the default method in Suite2p ^71^. The outcome of spike deconvolution was the inferred spike probability up to an unknown multiplicative constant independent for each neuron. We later normalized all neural responses at the end of the preprocessing stage, and thus, the unknown multiplicative constant was not influential.

#### Retinotopic mapping

We initially mapped the retinotopy before the adaptation experiments to determine where to place a stimulus in a given recording. To do this mapping, we used sparse noise stimuli, consisting of black or white squares with a width of 6 deg visual angle on a grey background, which were presented to the mouse for 30 min. Squares appeared randomly at fixed positions in a 15 by 45 grid spanning the retinotopic range of the computer screens. At any one time, 2% of the squares were shown.

#### Normalization of neural responses

For each neuron *i*, trial *t*, and environment *l*, spontaneous activity *x*_*s,i,l*_(*t*) was computed as the average inferred spike probability over 250 ms before the stimulus onset while evoked activity *x*_*e,i,l*_(*t*) was computed as the average inferred spike probability for the whole stimulus duration of 500 ms.

We then detrended the activity in the way described in this paragraph. We considered the average spontaneous activity, averaged across neurons *x̅*_*s,l*_(*t*) ≡ 〈*x*_*s,i,l*_(*t*)〉_*i*_, for each of the three environments (uniform #1, biased, uniform #2). We then separated the values of *x̅*_*s,l*_(*t*) for each trial depending on the running speed (stationary: v < 1cm/s; low speed: 1 cm/s < v < 15 cm/s; high speed: v > 15 cm/s). In each environment, we considered the locomotion condition out of these three (stationary, low speed, high speed) with the largest number of trials. We finally fit an exponential *f*_*l*_(*t*) = *ae*^−*bt*^ + *c* on the values *x̅*_*s,l*_(*t*) of this condition. We then divided *f*_*l*_(*t*) to all neurons from spontaneous *x*_*s,i,l*_(*t*) and evoked *x*_*e,i,l*_(*t*) by that in the given environment *l*, giving the new values *r*_*s,i,l*_(*t*) = *x*_*s,i,l*_(*t*)⁄*f*_*l*_ and *r*_*e,i,l*_(*t*) = *x*_*e,i,l*_(*t*)⁄*f*_*l*_ for spontaneous and evoked activity.

Finally, we z-scored the evoked activity as follows. For each neuron *i* and environment *l*, we computed the average spontaneous activity, averaged across environments *r̅*_*s,i*_ = 〈*r*_*s,i*_(*t*)〉_*t,l*_ and the standard deviation averaged across environments *σ*_*r,s,i*_ = 〈*σ*[*r*_*s,i,l*_(*t*)]_*t*_〉_*l*_. When then computed for each neuron *z*_*i,l*_(*t*) = [*r*_*e,i,l*_(*t*) − *r̅*_*s,i*_]⁄*σ*_*r,s,i*_.

#### Tuning parameters

For each neuron *i* and environment *l*, we computed the preferred orientation by first considering the average response of a neuron across repeats of the same stimuli in a uniform environment *z*_*i,l*_(*θ*) = 〈*z*_*i,l*_(*t*)〉_*t*∈*θ*_ to an orientation *θ*. We then computed *z̃*_*i,l*_(*θ*) = *z*_*i,l*_(*θ*) − min *z*_*i,l*_(*θ*). Then, the preferred orientation corresponded to the circular mean *φ*_*i*_ = 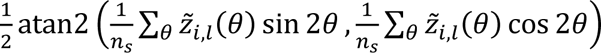, where *n*_*s*_ = 12 is the number of stimuli. To estimate average tuning curves based on preferred orientations (also known as super-neurons), we grouped each neuron into one of *n*_*s*_ groups based on the closer discrete orientation. We then took the average responses across neurons without normalizing their responses (other than the normalization already described in a previous session), giving rise to *z*_*e,l*_(*θ*, *φ*), where *φ* it the assigned preferred orientation of a super-neuron.

To compute a phenomenological model of the Euclidean distances between average tuning curves we proceeded as follows: we first fitted these average tuning curves to von Mises function and estimated the amplitude a_φ_, concentration parameter κ_φ_, and offset c_φ_ : *m*(*θ*, *φ*) = a_φ_*f*_*VM*_ (2*θ*; κ_φ_, 2*φ*) + c_φ_, where *f*_*VM*_ is the von Mises density function *f*_*VM*_ (*x*; κ, *µ*)= exp[κ cos(*x* − *µ*)]⁄[2*πI*_0_ (*κ*)] and where *I*_0_(*κ*) is the modified Bessel function of the first kind of order 0 (we dropped the environment index *l* as this analysis was done only in the uniform environment). Then, we computed an average concentration parameter κ̅, together with an average amplitude a̅, and an average offset c̅, by fitting a single von Mises function *m̅*(*θ*, *φ*) = c̅*f*_*VM*_(2*θ*; κ̅, 2*φ*) + c̅ after averaging all individual responses, which were circularly shifted based on their preferred orientation. After this, we considered two cases: (i) fixed concentration but variable amplitudes or (ii) fixed amplitude but variable width.

We also fitted a von Mises function to the tuning curve of each individual neuron in the uniform environments, a_i_*f*_*VM*_(2*θ*; κ_i_, 2*φ*) + c_i_ and took the concentration parameter κ_i_ as a measure of tuning sharpness. For each bin of preferred orientation we report the median of κ_i_ across all neurons (Fig. 1e) or across the neurons whose fit explained more than 75% of the variance (R^2^ > 0.75, Fig. S1e); the same quantity computed on super-neurons is shown in Fig. S1f. We quantified orientation selectivity with the global orientation selectivity index, *gOSI*_*i*_ = 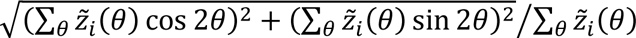, and considered a neuron tuned when gOSI_i_≥0.4. Super-neurons were computed from tuned neurons, except in Fig. S4a,b, where all neurons were included.

#### Alignment to the adaptor

Matrices of responses, distances and discrimination accuracies are displayed as a function of the distance from the adaptor. We circularly shifted each matrix so that the adaptor fell at its center, and averaged the shifted matrix with its reflection about the adaptor, which exploits the symmetry of the stimulus set around the adaptor orientation.

#### Distances between population responses

For each stimulus θ, environment and locomotion condition, we computed the mean response vector r(θ) across trials. We measured the separation between the representations of two stimuli either by their Euclidean distance ‖r(θ_i_)-r_j_(θ)‖ or by their cosine distance, d(θ_i_,θ_j_)=1-(r(θ_i_)·r(θ_j_))/(‖r(θ_i_)‖‖r(θ_j_)‖). Trial-to-trial variability biases the cosine distance upwards, so we compared it only between conditions with matched numbers of trials. We also expressed the Euclidean distance in units of the response variability, dividing it by the standard deviation across repeats averaged over neurons and stimuli within the same environment, d_n_ (θ_i_,θ_j_)=‖r(θ_i_)-r(θ_j_)‖/⟨σ⟩ (Fig. S4c,d).

#### Visualization of the population geometry

To visualize the geometry, we projected the responses onto their first two or three principal components. For the model, we rotated the projection so that the adaptor lay on the vertical axis and translated it so that the response to the blank stimulus lay at the origin (Fig. 4c, S14a, S15a-c, S16a).

#### Response variability

For each neuron, stimulus and environment we computed the standard deviation σ of the responses across repeats of the same stimulus. To test whether changes in variability contributed to the changes in discriminability, we rescaled the responses of each neuron so that σ took a single value *σ*_0_, common to all stimuli and to both environments, while preserving the mean response *m* to each stimulus, *z*′_*i,l*_(*t*) = *z*_*i,l*_(*θ*) + [*z*_*i,l*_(*t*) − *z*_*i,l*_(*θ*)]*σ*_0_⁄*σ*_*i,l*_(*θ*), and we repeated the decoding analysis (Fig. S5c,d).

#### Noise correlations

We measured noise correlations as the Pearson correlation between the responses of pairs of neurons across repeats of the same stimulus, after subtracting the mean response to that stimulus and equalizing the number of trials across environments. To remove correlations we shuffled the trials of each neuron independently, separately for each stimulus and environment, and repeated the decoding analysis (Fig. S2b-d, S5g,h).

#### Discriminability

We decoded the orientation of the static gratings presented to the mice using the population activity from V1 neurons. Let us consider environments with different stimulus statistics, i.e. different probability of stimulus presentation with a given orientation. For each environment, we computed the discrimination accuracy between any pair of stimuli. When considering the same environment, we equalized the number of trials for each recording session in the following way: we computed the minimum number of repeats for a given orientation across all 12 orientations. Then, we used 2/3 of these repeats for training the model and 1/3 for testing the model. We used a Linear Support Vector Classification as a model trained on all neurons recorded for a particular session with a regularization parameter equal to 0.1. We then tested the discrimination accuracy of the model on the test data.

In other analyses, training and testing data were from different environments (e.g., training in a uniform condition and testing in a biased condition). We still computed the minimum number of repeats per stimulus in the training and testing environment separately in those cases. We did not need to take only a fraction of this minimum number.

We also computed discrimination accuracy in low-dimensional subspaces, by projecting the responses onto their first principal components, computed on all the trials of the uniform environments, and training the same classifier on an increasing number of components (Fig. S6).

To separate the effect of the environment from that of the passage of time, we compared decoders trained and tested in different time blocks. We measured the discrimination accuracy between the responses to the same stimulus in two blocks, either two uniform environments or a uniform and a biased environment, as a function of the number of blocks separating them (Fig. S8c-e), and we repeated the analysis of Fig. 2i training the decoder in the uniform environment adjacent to both test environments (Fig. S8a,b).

#### Cross-condition generalization performance

To compute the cross-condition generalization performance (CCGP) ^15,49^, we typically considered two different variables, for example, stimulus orientation *θ* and locomotion condition (stationary and running). We then trained, for example, a linear decoder to discriminate between two angles, *θ*_1_ and *θ*_2_, during the stationary condition and tested the decoder to discriminate the same angles in the running condition. We sampled the number of trials to balance them across classes separately in the training and test set.

We applied the same procedure to other pairs of variables. To test whether the coding directions for adaptation were preserved across locomotion states, we trained the decoder to discriminate the uniform from the biased environment in one locomotion condition and tested it in the other, and compared the result with the discrimination of two uniform environments (Fig. S12c,d); conversely, we trained the decoder to discriminate the stationary from the running condition in one environment and tested it in another (Fig. S12e,f). We also trained the decoder to discriminate stationary from running trials at one orientation and tested it at another orientation, with preserved or shuffled trials (Fig. S11d-f).

#### Time course of adaptation

To compare the two uniform environments we divided the responses of each block by their mean across neurons and stimuli and we subtracted from each session the average change in discrimination accuracy across all pairs of stimuli (Fig. S9b). To follow the changes within the biased environment we recomputed the mean responses and the cosine distances in sliding windows of 200 trials, in steps of 20 trials (Fig. S9d,e).

#### Locomotion

We classified a trial as running when the average running speed during the stimulus exceeded 0.5 cm/s, and as stationary otherwise. A recording session contributed to a locomotion condition when it had enough trials of every orientation to train and test the decoder in that condition, so that sessions were selected independently for the two conditions. We compared the changes in response, discrimination accuracy and cosine distance between the two conditions with two-sided Mann-Whitney U tests (Fig. 3j-l).

#### Scaling model

We asked whether locomotion rescales the population responses, and described the change from the stationary to the running condition as a truncated cone. Let x(θ) and y(θ) be the mean responses to stimulus θ during stationary and during running periods, averaged over the two uniform environments, and let *m*_*stat*_ and *m*_*run*_ be their averages across stimuli. We placed the apex of the cone on the line joining the two, *x*_0_ = *m*_*stat*_ + *a*(*m*_*run*_ − *m*_*stat*_), and modeled the running response to each stimulus as an expansion of the stationary response about that apex, *x*^(*θ*) = *x*_0_ + *g*[*x*(*θ*) − *x*_0_]. We estimated the position of the apex a and the gain *g* on a grid by minimizing the normalized error *E* = 〈‖*x*^(*θ*) − *y*(*θ*)‖^2^〉⁄〈‖*x*(*θ*) − *y*(*θ*)‖^2^〉, where the averages are taken over the 12 stimuli, so that *E* = 1 corresponds to a model that leaves the responses unchanged. We report *E* averaged across the sessions with enough stationary and running trials for every stimulus (Fig. S11g).

#### Angles between directions in the activity space

For each stimulus we defined the direction of adaptation as the difference between the mean response vectors in the biased and in the uniform environment, the direction of locomotion as the difference between the running and the stationary condition, the stimulus-coding direction as the difference between the mean responses to adjacent orientations, and the direction of maximum variability as the first principal component of the responses to that stimulus across trials. We report the cosine of the angle between pairs of these directions (Fig. S11b,c, S12a,b).

#### Statistics

Statistical tests were two-sided and computed across recording sessions. We used a one-sample t-test for changes in discrimination accuracy, a Wilcoxon signed-rank test for changes in responses, distances and correlations, and a Mann-Whitney U test to compare the stationary and the running condition.

### Normative Model

#### Model training

We trained an autoencoder model with three different layers: an input layer ***x*** with *n*_*x*_ = 121 units (corresponding to a vectorized 11×11 image), a hidden layer ***r*** with *n*_*r*_ units, an output layer ***y*** with *n*_*y*_ = *n*_*x*_ units. We used sigmoid activation functions. The goal of the autoencoder was to minimize a cost function *E* = *E*_1_ + *λE*_2_ where 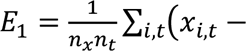 *y_i,t_*)^2^ and 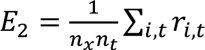, which, since *r*_*i,t*_ > 0, represents an L1 sparsity constraint (metabolic cost). We added Gaussian noise with strengths *ε*_*x*_ and *ε*_*r*_ = 1.0 in the input and hidden layers. We trained the autoencoder using Pytorch using a batch size of 500 and a max number of epochs of 3,000 if convergence was not reached. We set the convergence error to 0.05 · 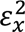. We implemented gradient descent with the Adam algorithm with a learning rate of 10^-4^.

We scanned through three different hyperparameters: the penalty factor on the metabolic cost *λ* ∈ {0.,0.1,0.25,0.5,1.}, the noise in the input *ε*_*x*_ ∈ {0.25,0.5,1.,2.}, the number of hidden units *n*_*r*_ ∈ {500, 1,000, 2,000, 3,000, 5,000} for a total of 100 of different sets of hyperparameters. We repeated 30 trainings with random initializations for each of these parameters. For the analysis of small network size we trained additional networks with *n*_*r*_ ∈ {24,96,384} hidden units, with three random initializations each (Fig. S15a-c).

#### Stimuli presented to the model

Similarly to the experiments, we presented sequences of oriented gratings plus blank stimuli with zero contrast to the model. Both uniform and biased environments had 2,000 blank stimuli. Then, 120 trials were added with 10 repeats of one orientation for each of the 12 considered orientations. In addition, 1,000 other stimuli were drawn from a von Mises distribution with a concentration parameter of 0 in the uniform environment and a concentration parameter of 150 in the biased environment. Stimuli had a size of 11×11 pixels and corresponded to a Gabor function with a spatial scale of 10 pixels and spatial frequency of 1.5 pixels. To test the contribution of the blank stimuli we also trained networks in which the number of blanks varied from 0 to 2,000 while the total number of presentations was held at 3,000 (Fig. S14).

#### Decoding of model responses

We computed discrimination accuracy using the same method used for the neural data but we only used a subset *n*_*d*_ = 50 of the artificial neurons to decode the direction of motion from the population activity. In contrast to the training procedure, for the decoder we used the trained models as forward models but changed the level of noise in the input to 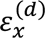 = 0.3 and, similarly, in the hidden layer to 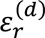 = 0.2. The values of *n_d_*, 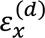, and 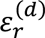, where chosen to ensure compatibility on the experimental results on discrimination accuracy.

#### Sparsity of the model

We quantified the sparsity of the hidden layer with the Gini index of the responses across hidden units, 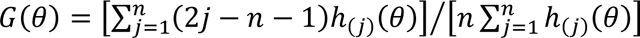, where ℎ_(1)_(*θ*) ≤ ⋯ ≤ ℎ_(*n*_*r*_)_(*θ*) are the responses to one stimulus sorted in increasing order, so that higher values indicate sparser responses. We report the index averaged over the 12 gratings (Fig. S14d, S15d,e).

#### Comparison between model and data

We compared each model with the data by correlating the model and the experimental matrices of the difference in discrimination accuracy between environments, across all pairs of stimuli, and the corresponding matrices of the difference in normalized responses, across preferred orientations and stimuli. Each point in Fig. 4i is the average across the 30 networks trained with the same hyperparameters.

#### Convolutional autoencoder

We repeated the analysis with a convolutional autoencoder, whose encoder was a convolutional layer with 5×5 filters applied with a stride of 3, giving 3×3 spatial positions for each channel, and whose decoder was the corresponding transposed convolution. The cost function, the noise, the stimuli and the decoding were as in the fully connected model. The network shown had 240 channels and 12,241 parameters, 1.7% of those of the fully connected model with *n*_*r*_ = 3,000, and the results were averaged over 30 trained networks (Fig. S16).

## Supplementary Figures

**Supplementary Figure 1.**
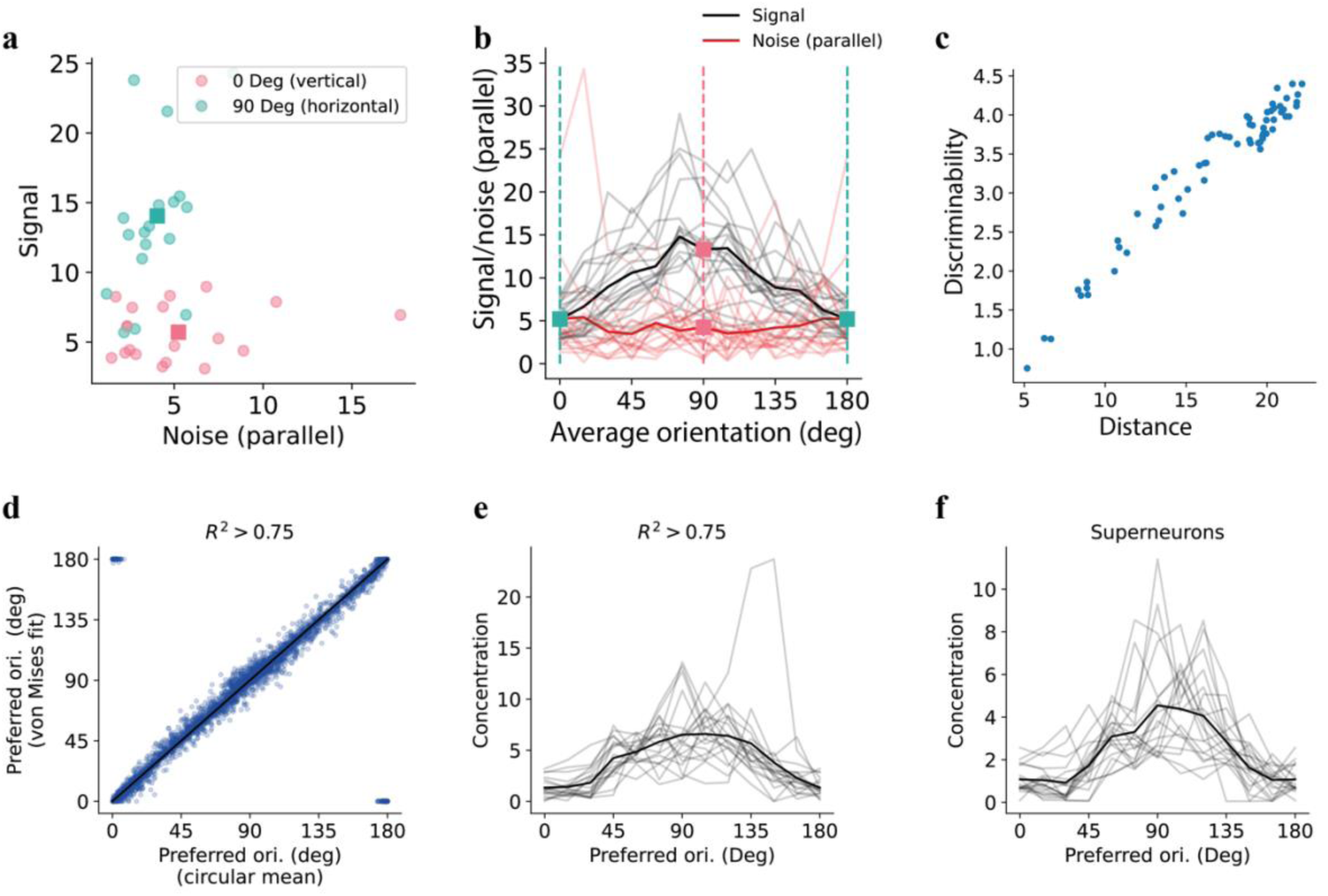
related to Figure 1. Differences in discriminability across orientations reflect primarily differences in the signal (response distance). **a)** the y-axis is the “signal” i.e. the Euclidean distance between the average neural response vectors of two different stimuli in the uniform environment; the x-axis is the “noise” or the vector of average standard deviation projected parallel to the signal; one of the two stimuli is either the horizontal (green, 90 deg) or vertical (red, 0 deg) orientation and the other is a stimulus 15 deg away; each point is a different recording session, squares are averages across all recording sessions; **b)** similar to (a) but signal (black solid lines) and noise (red solid lines) are plotted separately as a function of the average angle of the pairs of orientation considered. Thin lines are individual recording sessions; thick lines are average across recordings. For reference, the vertical dashed lines correspond to the average angles in (a); **c)** discriminability between different pairs of orientations vs. normalized distances, averaged across experimental sessions. **d)** Preferred orientation estimated as circular mean or through a fitting of tuning curves with von Mises distributions. **e)** Average concentration parameter of neurons with a good von Mises fit (R^2^ > 0.75) as a function of their preferred orientation. Gray lines are individual recording sessions, while the black line is the average across sessions. **f)** Same as (e) but for superneurons.

**Supplementary Figure 2.**
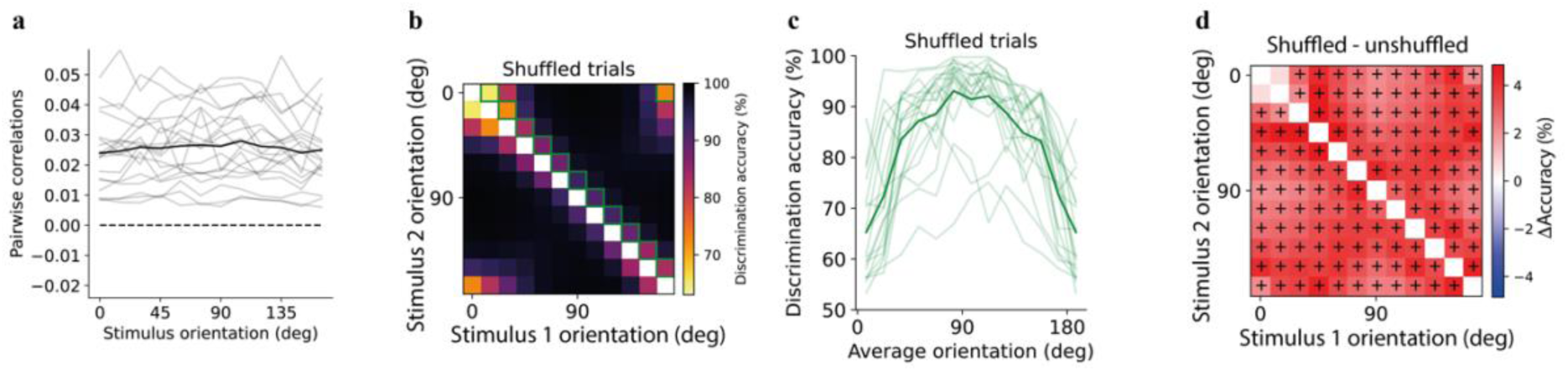
related to Figure 1. Noise correlations modestly reduce, but do not reshape, orientation discriminability. **a**) Average pairwise correlations in the uniform environment as a function of stimulus orientation. **b,c**) Same as Fig. 1i,j but shuffling the trials for each stimulus. **d**) Difference in discrimination accuracy between decoder with destroyed (b,c) or preserved (Fig. 1i,j) correlations; pluses correspond to a significant increase in discrimination with destroyed correlations (p < 0.05, Wilcoxon signed-ranked test).

**Supplementary Figure 3.**
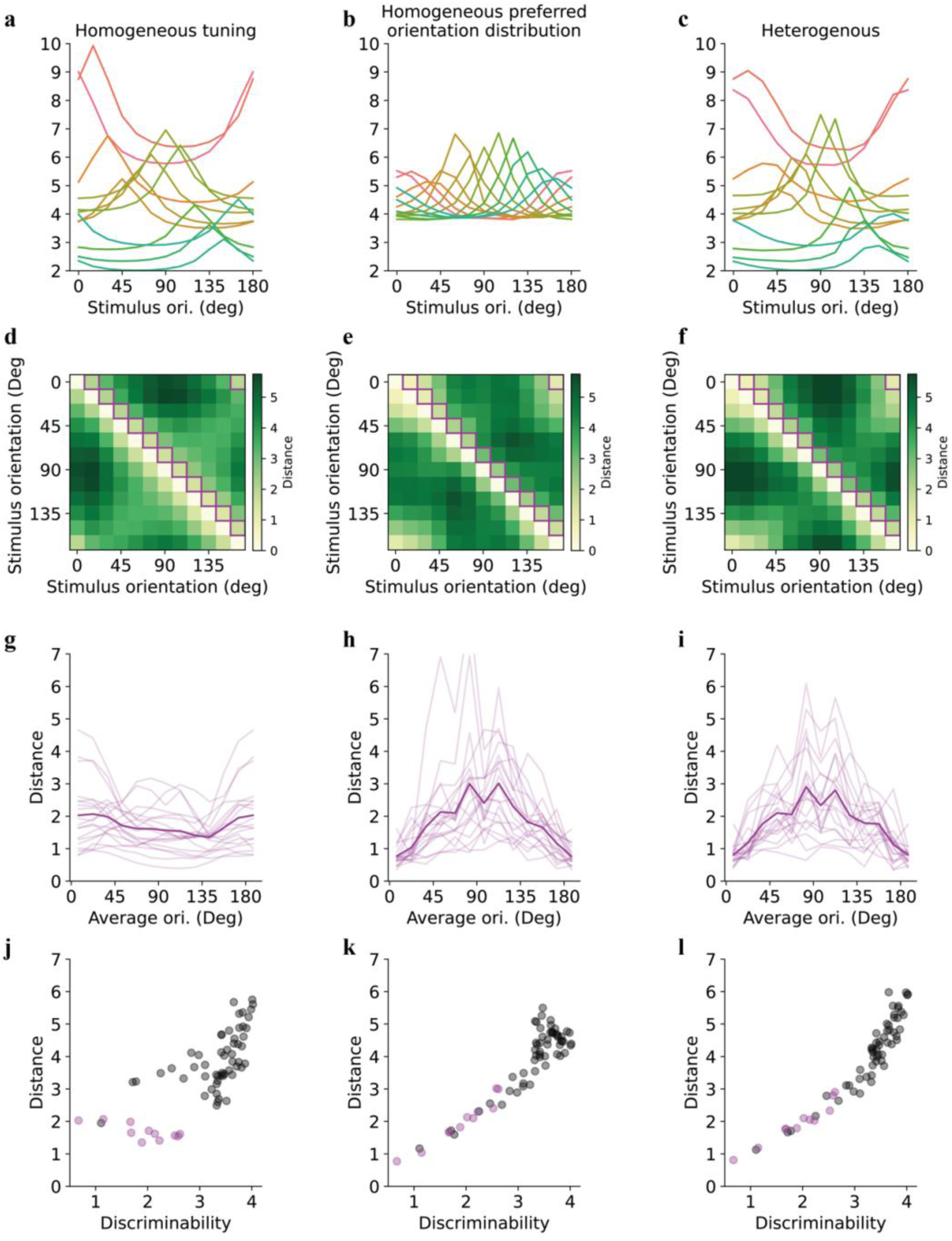
related to Figure 1. Both the non-uniform distribution of preferred orientations and of the tuning bandwidth shape discriminability. **a**) Average responses of neurons with similar orientation preferences after taking into account the difference in the distribution of orientation preferences (Fig. 1d) but not the tuning bandwidth (Fig. 1e): tuning curves have a fixed bandwidth but response strength proportional to the distribution of orientation preference. **b**) Average tuning curves of neurons with variable bandwidth but assuming a fixed distribution of orientation preference. **c**) Average tuning curves with variable bandwidth and distribution of orientation preferences. **d-f**) Normalized distance between stimuli based on the tuning curves in (a-c), plotted by assuming one neuron with each tuning curve in (a-c) and homogeneous noise in the responses. **g-i**) normalized distances between nearby stimuli (separated by 15 deg) corresponding to purple squares in (d-f); **j-l**) Euclidean distances between pairs of stimuli vs. discriminability; purple circles correspond to pairs reported in (g-i).

**Supplementary Figure 4.**
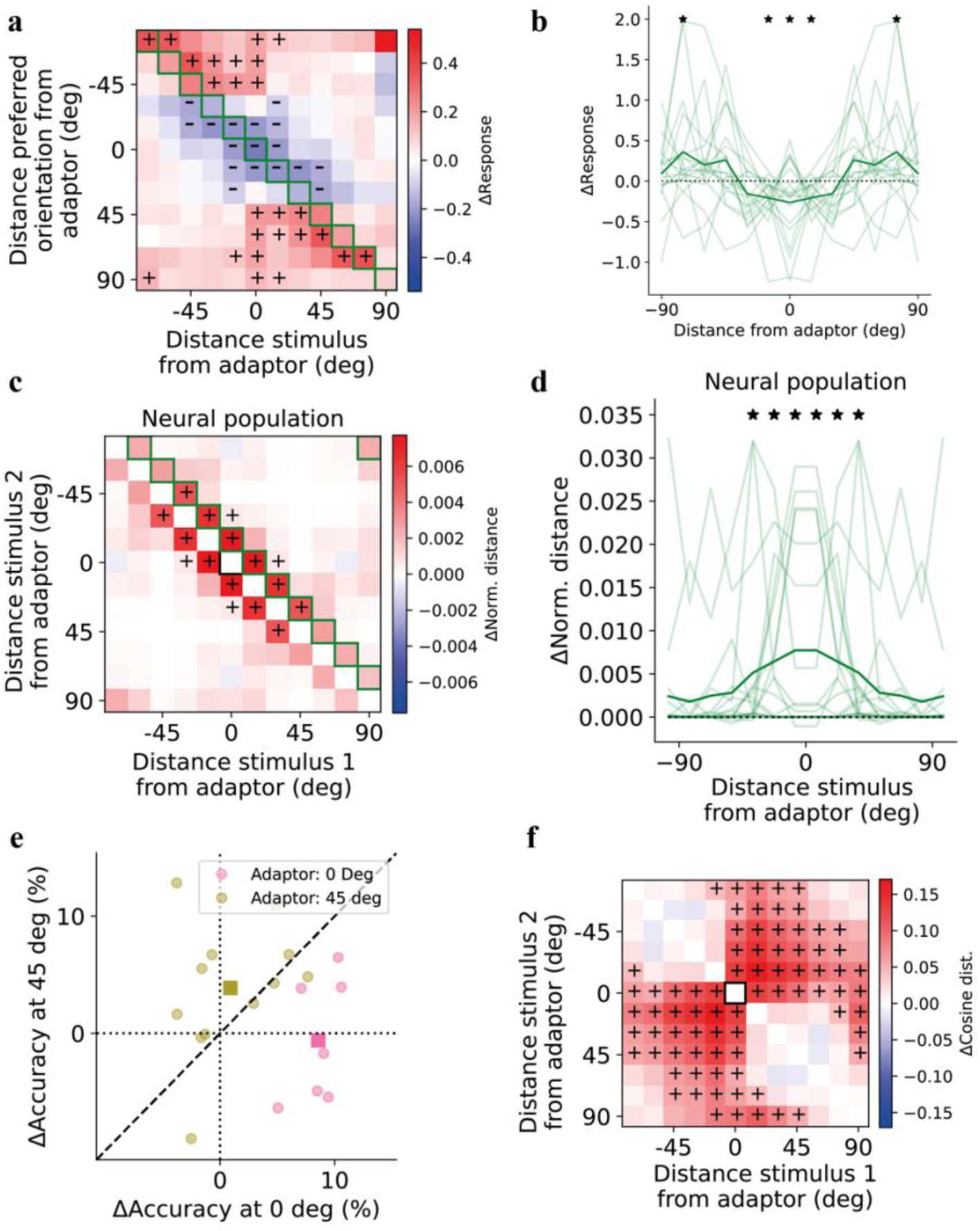
related to Figure 2. The expansion of representations near the adaptor is robust to tuning-strength criteria and distance metrics. **a,b**) Same as Fig. 2e,f but also including neurons with low orientation tuning (gOSI < 0.4). **c**) Difference in normalized Euclidean distance in neural space between any pair of orientations averaged across sessions; green squares correspond to average values in the right panel; pluses correspond to a significant increase in discrimination (p < 0.05, 1-sample t-test). **d**) **C**hange in normalized Euclidean distance between biased and uniform environments for stimuli 15 deg apart as a function of the distance of the stimuli from the adapter; thin lines correspond to each recording session, thick line is an average across all sessions; asterisks are a significant increase in normalized Euclidean distance. **e**) Change in discrimination accuracy from uniform to biased conditions in discriminating orientation near the adaptor or 45 deg from it. Each dot corresponds to a recording session, squares correspond to the average across sessions where the adapter was at 0 deg (vertical orientation) or 45 deg (oblique orientation). **f**) Change in cosine distance between angles (corresponding to 1 − cos *α*, where *α* is the angle between two vectors, thus higher cosine distance corresponds to larger angles) from a uniform and a biased environment; black square corresponds to adapter orientation; pluses correspond to a significant increase in discrimination (p < 0.05, 1-sample t-test); note that except for pairs of orientations near the adaptor significant increases in cosine distances for a pair of angles do not necessarily correspond to a significant increase in Euclidean distances or discrimination accuracy; this is because the three measures are not exactly the same in the way the account for noise and other factors.

**Supplementary Figure 5.**
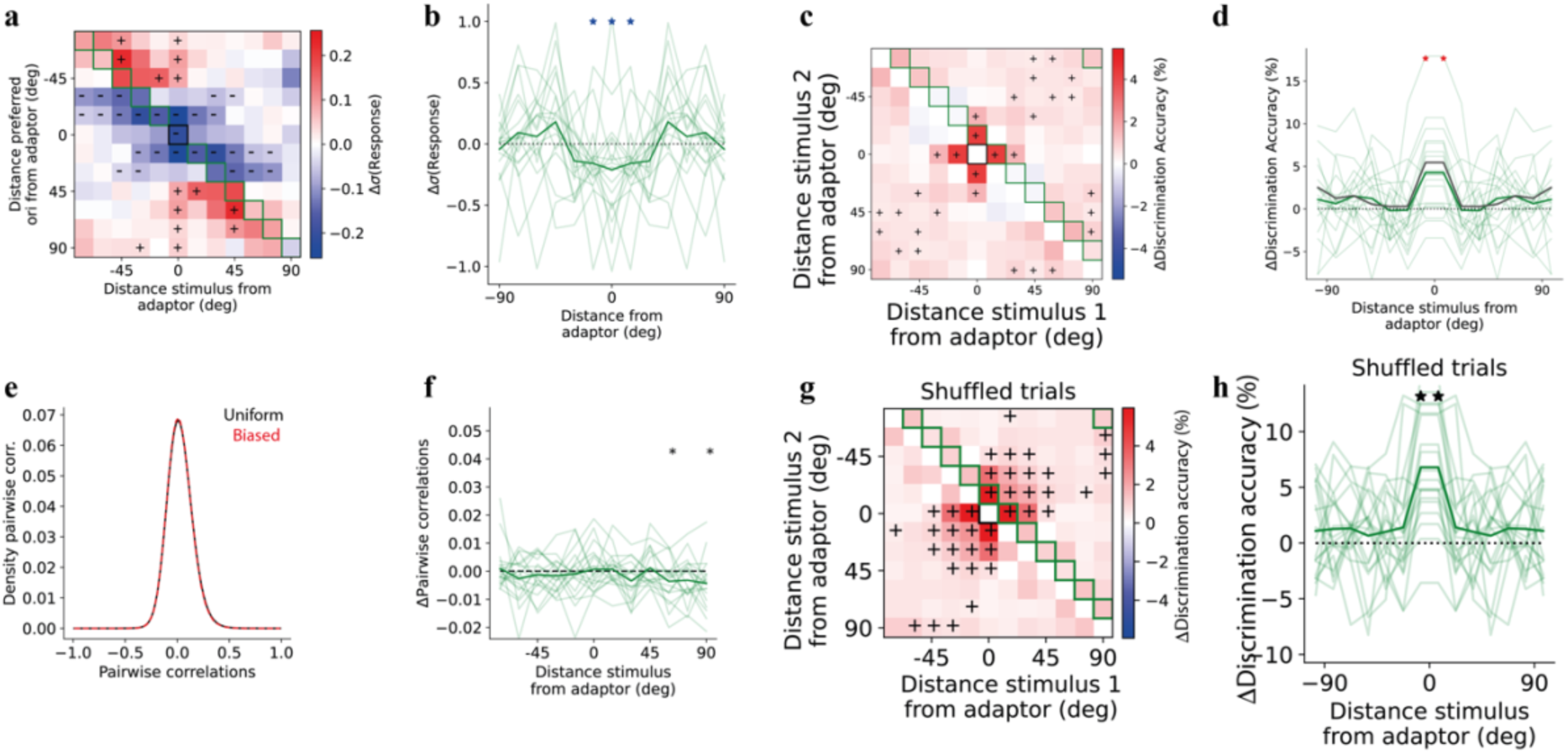
related to Figure 2. Response variability and correlations are not sufficient to explain discriminability increase near the adaptor. **a)** Change in response variability (standard deviation, σ) between biased and uniform environments averaged across recording sessions, similar to Fig. 2e but on per-cell σ instead of mean response; minuses correspond to a significant decrease in variability (p < 0.05, 1-sample Wilcoxon); green squares correspond to values in (b). **b)** Δσ at the preferred orientation as a function of distance from the adapter; light green: individual recording sessions; dark green: average across sessions; blue asterisks: significant decrease (p < 0.05). **c)** Same as Fig. 2i but on data with per-stimulus σ equalized across environments (per-stimulus means preserved); color scale matches Fig. 2i; pluses: significant increase (p < 0.05, 1-sample t-test); green squares correspond to values in (d). **d)** Same as Fig. 2j but for the variance-equalized SVM (green) overlaid with the control SVM mean (dark grey; equivalent to the thick line in Fig. 2i); light green: individual sessions; asterisks: significant increase (p < 0.05). **e)** Distribution of pairwise correlations in uniform and biased environments. **f**) Difference in average pairwise correlations between biased and uniform environments as a function of the distance of the stimulus from the adaptor. **g,h**) Same as Fig. 2i,j but shuffling the trials for each stimulus.

**Supplementary Figure 6.**
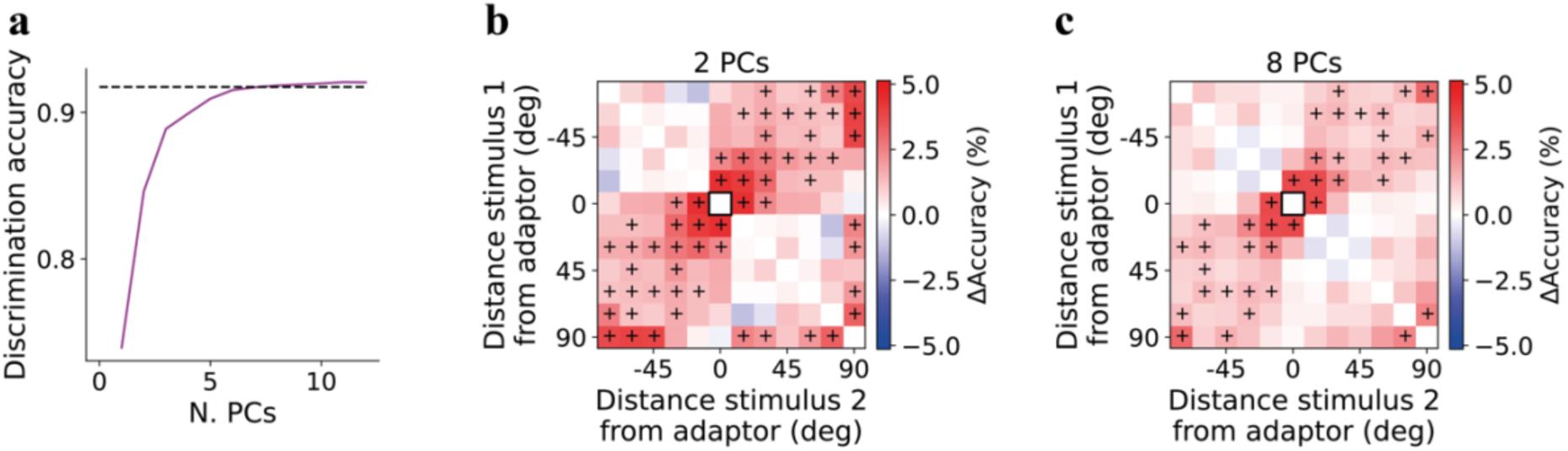
related to Figure 2. Adaptation increases discriminability near the adaptor even in low-dimensional subspaces. **a**): average discrimination accuracy as a function of the number of PCs considered. **b**): change in discriminability when using only the first two PCs. **c**): change in discriminability when using the first eight PCs.

**Supplementary Figure 7.**
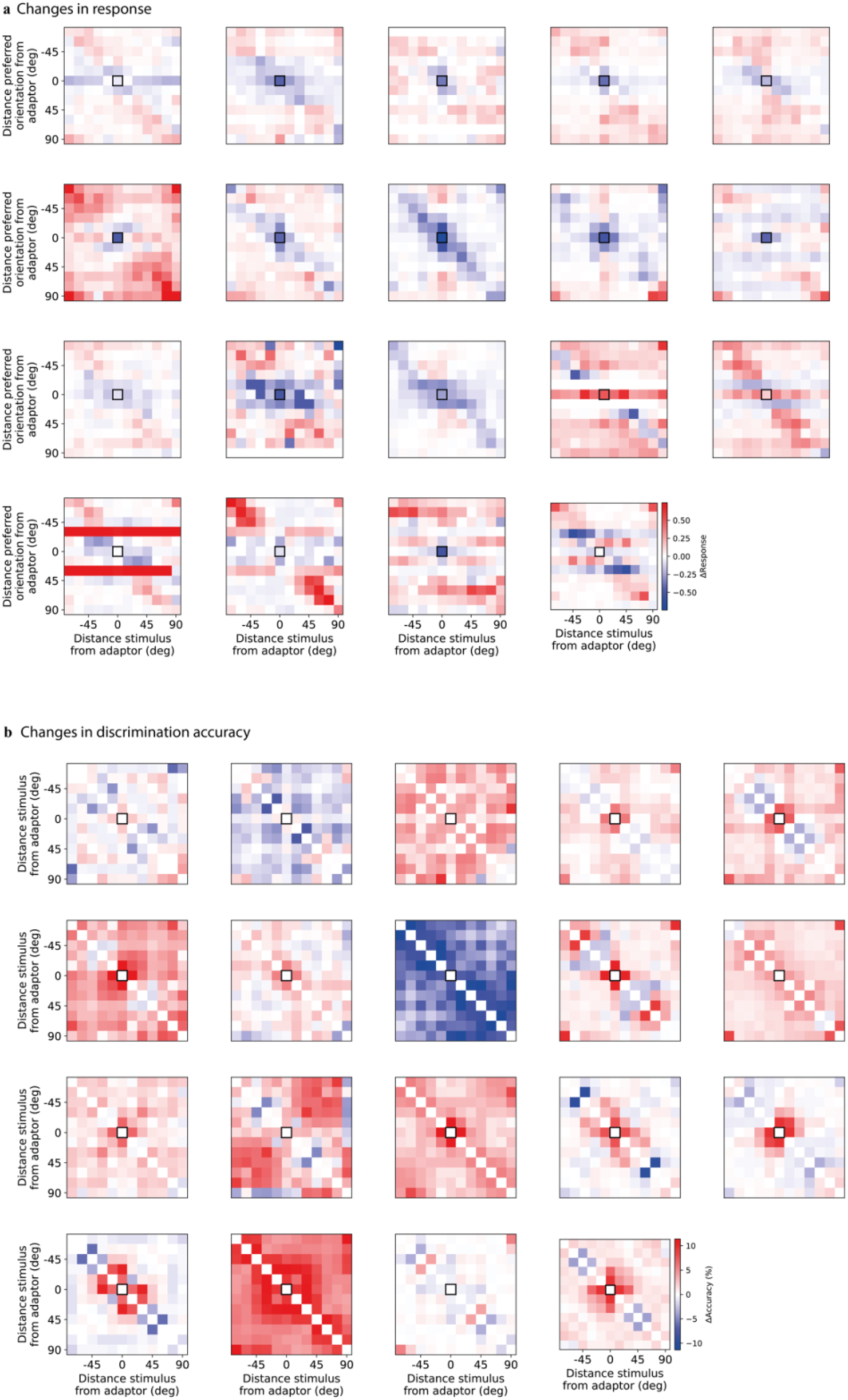
related to Figure 2. Response and discriminability changes near the adaptor are largely consistent across recording sessions. **a)** Same as Fig. 2d but for all recording sessions and after applying transformations reflecting symmetries around the adaptor; white horizontal stripes indicate the absence of neurons at that preferred orientation bin. **b)** Same as Fig. 2h but for all recording sessions and after applying transformations reflecting symmetries around the adaptor.

**Supplementary Figure 8.**
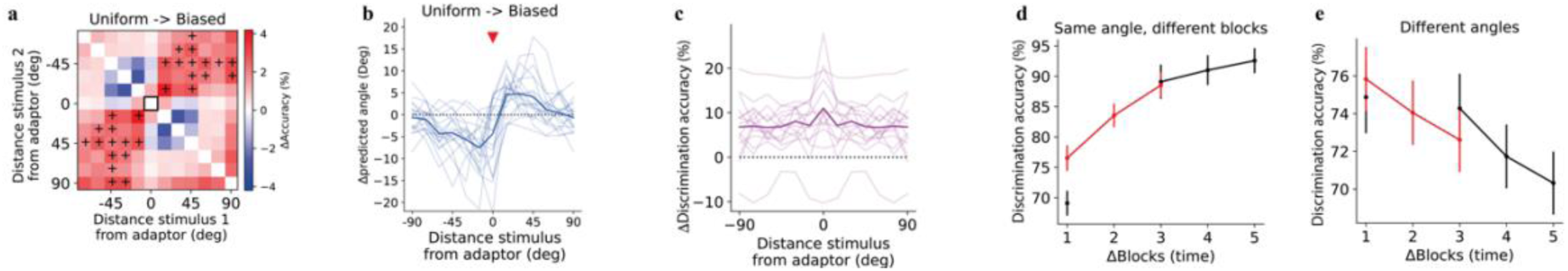
related to Figure 2. The increased discriminability is specific to the biased environment, not explained by time or block effects. **a**) Difference in discrimination accuracy between a biased and a uniform environment when the model was trained in a uniform environment temporally adjacent to the two test environments, averaged across experimental sessions. Pluses correspond to a significant increase in discrimination accuracy. **b**) Angle prediction in a biased environment after training a model in a uniform environment. The red triangle corresponds to the adapter’s relative orientation. **c**) Difference in accuracy to discriminate different time blocks, computed by subtracting discrimination accuracy when training and testing in the same environment (uniform) but in different (and adjacent) time blocks from the discrimination accuracy in the different environments (uniform and biased) and adjacent time blocks. Discrimination accuracy was computed for responses to the same stimulus each time and then plotted as a function of stimulus orientation; thin lines are values in each recording session (note one outlier recording session out of 19 sessions with differences below 0); the thick line is an average over all sessions. **d**) Discrimination accuracy between a stimulus in a time block and the same stimulus in another time block as a function of the number of experimental blocks separating the two sequences of stimuli (same as in c but for different time blocks and averaged across stimulus orientations). This is similar to (c), the difference being that time blocks are not only adjacent, and instead of the difference between discrimination accuracy in the same environments vs. different environments, the two are plotted separately: black corresponds to the average between training in a uniform environment and testing in another, while red corresponds to training in a uniform environment and testing in a biased environment. Error bars are computed across all experiments; **e**) same as in (d), but discrimination is between two different angles.

**Supplementary Figure 9.**
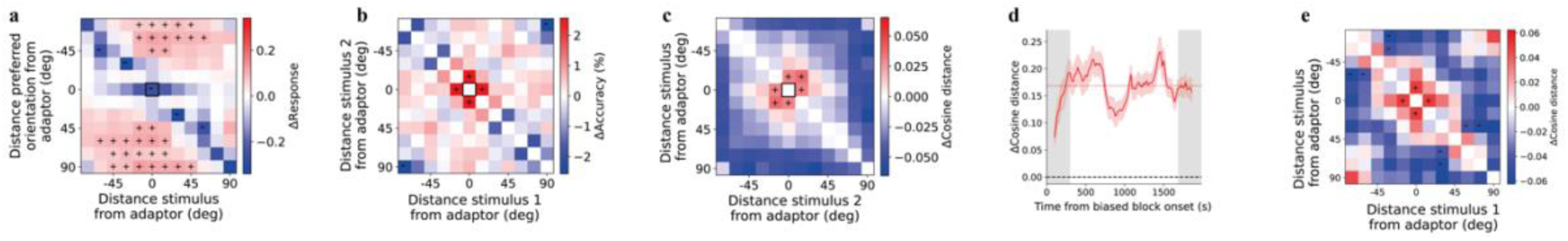
related to Figure 2. Adaptation-induced changes in geometry build up gradually and persist beyond the adapting period. **a)** The difference between the third block (U2) and the first block (U1), both uniform, averaged across sessions. Difference in normalized responses of tuned neurons at a given preferred-orientation distance from the adaptor (y-axis) to stimuli a given distance from it (x-axis), after normalizing each block by its mean response; black square: adaptor response of adaptor-tuned neurons; pluses (resp. minuses): significant increase (resp. decrease) (p < 0.05, Wilcoxon). As in Fig. 2e, neurons tuned near the adaptor decrease their responses to stimuli near the adaptor, and those tuned far from it increase their responses. **b)** Difference in discrimination accuracy between stimulus pairs at given distances from the adaptor, after subtracting each session’s mean change; pluses: significant increase (p < 0.05, one-sample t-test). Discriminability stays elevated near the adaptor, as in Fig. 2i. **c)** Difference in cosine distance between stimulus representations, pluses (resp. minuses): significant change (p < 0.05, Wilcoxon). Representations near the adaptor remain more separated. **d**) Build-up of representational geometry within the biased block: Cosine distance between the adaptor and the stimulus 15° from it, relative to its uniform (first-block) baseline, over the biased block (200-trial sliding windows, 20-trial steps). Red line: mean across sessions; shading: SEM; red dotted line: biased-block average; gray bands: the first-300 and last-300-trial windows differenced in (e).**e)** Change in cosine distance between pairs of stimuli within the biased block (last 300 trials minus first 300 trials, the shaded areas in d), as a function of each stimulus’s distance from the adaptor. Pluses (minuses): significant increase (decrease) across the 18 recording sessions (p < 0.05, Wilcoxon signed-rank test).

**Supplementary Figure 10.**
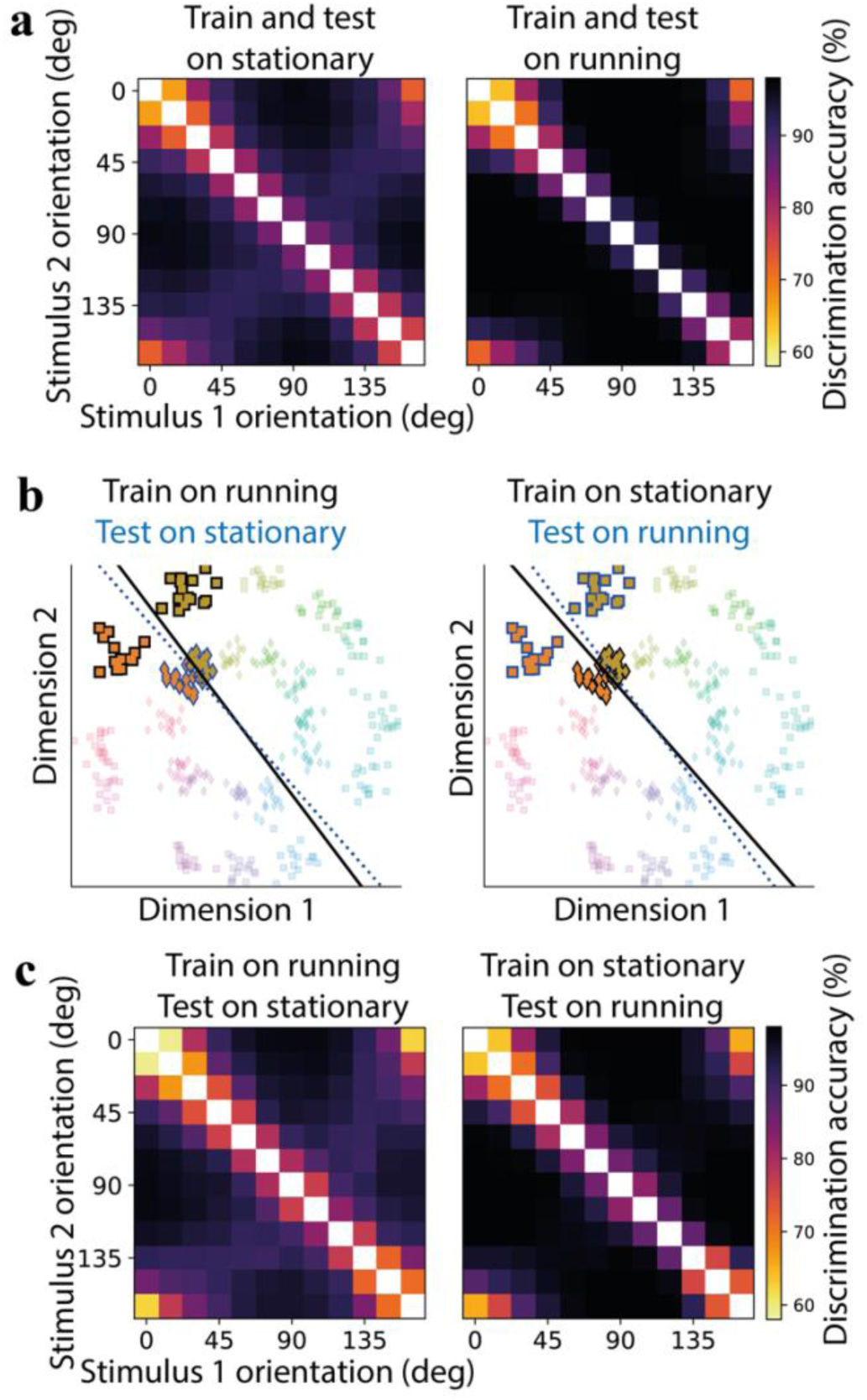
related to Figure 3. Running expands the geometry of representations. **a**) Discrimination accuracy between pairs of orientations for a model that has been trained and tested during stationary (left) or running (right) periods; **b**) Cartoon illustrating the measurement of cross-condition generalization performance (CCGP); left: a model was trained (black contour and black line) to discriminate two orientations (more opaque colors) during the running condition (squares) and then tested (blue contours) to discriminate the same orientation during the stationary condition (diamonds); for illustration purposes, we also plotted the hyperplane separating the points trained in the other condition (blue dotted line); right: same as before but model was trained in the stationary condition and tested in the running condition. **c**) Discrimination accuracy between pairs of orientations during running for a model that has been trained during stationary periods (left) or vice versa (right).

**Supplementary Figure 11.**
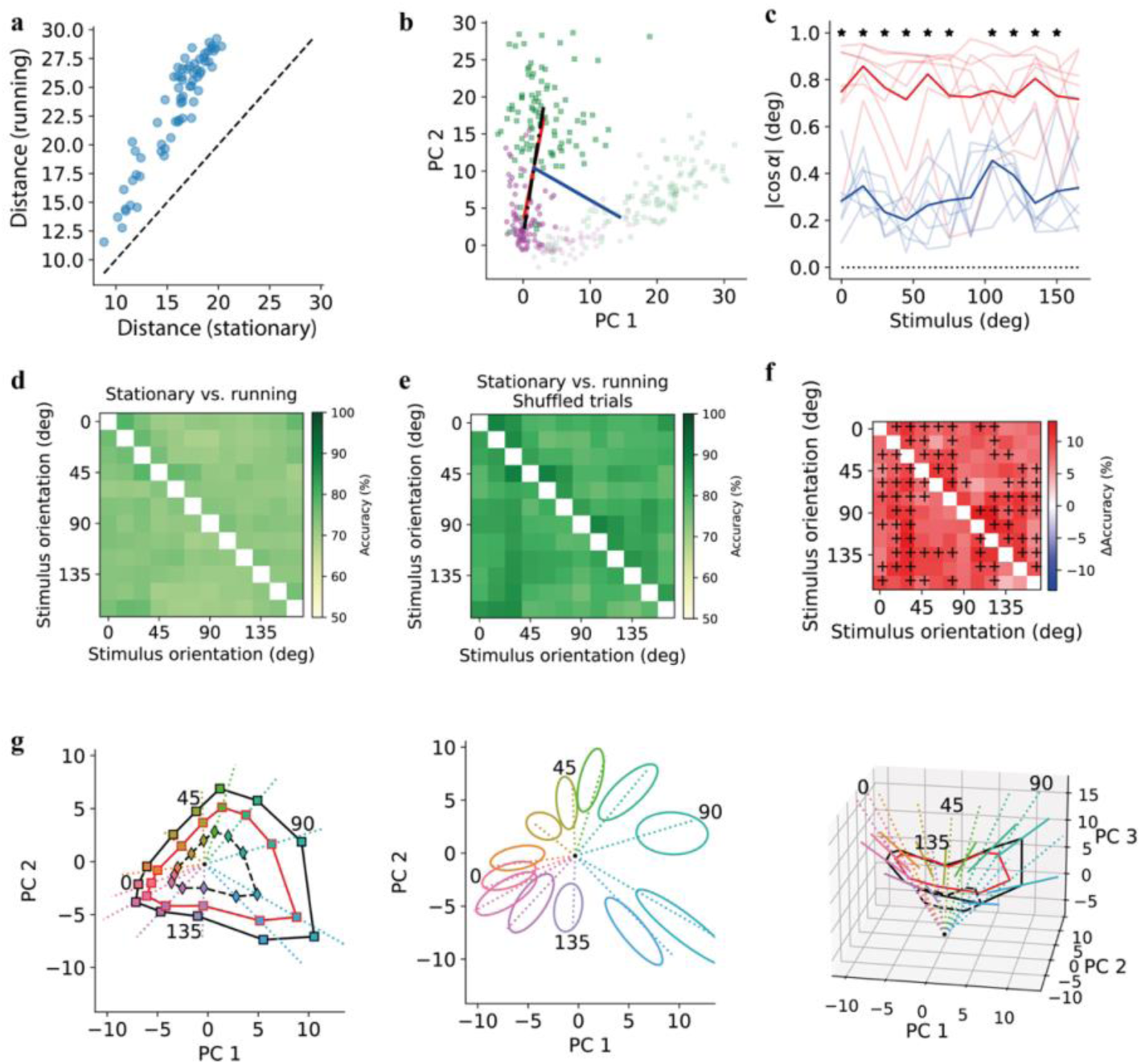
related to Figure 3. Locomotion modulates responses along the principal axis of variability, separate from stimulus-coding directions. **a**) Average Euclidean distance across experiments between any pair of stimuli; **b**) Representation of vectors to compute angles between stimulus (solid blue line), running (solid red line), and main axis of variability directions (dashed black line); green squares: running trials, magenta circles: stationary trials; transparency level indicates either of two different orientations. **c**) Blue lines: the cosine angle between the response vector of the first principal component of trials for each stimulus in neural space and the vector between adjacent stimuli; red lines: the cosine angle between the same first vector and the vector formed by stationary and running trials (running vector). Each thin line is one of the recording sessions where there was a sufficient number of stationary and running trials for each stimulus, the thick line is an average across sessions; **d**) Discrimination accuracy for a model that has been trained to discriminate stationary and running trials in an angle (x-axis) and tested in another angle (y-axis). **e**) Discrimination accuracy for a model that has been trained to discriminate stationary and running trials in an angle (x-axis) and tested in another angle (y-axis), same as (d) but shuffling the trials for each stimulus. **f**) Difference in discrimination accuracy for models that have been trained to discriminate stationary and running trials with preserved correlations (d) and models where correlations were destroyed (e); pluses correspond to a significant increase in discrimination (p < 0.05, Wilcoxon signed-ranked test). **g**) Left: first two components of a PCA on neural population responses plotted together with an affine model of responses during running (solid red line) based on scaling and shifting responses during stationary periods. Diamonds (resp. squares) are average stimulus responses during stationary (resp. running) period; the black line connects similar stimuli; the dotted colored lines connect stationary responses to the modeled running responses and represent the scaling with an offset used to obtain the model; center: ellipses are centered in the average responses during running in (g,left); the ellipses’ axes are proportional to the square root of eigenvalues of the stimulus-conditioned covariance matrix of the trial responses during running in the projected space; the dotted lines correspond to the same lines in (g,left); right: similar to (g,left) visualizing one extra PC; lines are centered in the average responses and their lengths correspond to the main axis of the ellipsoid in (g,center) but in three dimensions; the solid red line and the dotted lines are the same as in (g,left) but considering the extra PC.

**Supplementary Figure 12.**
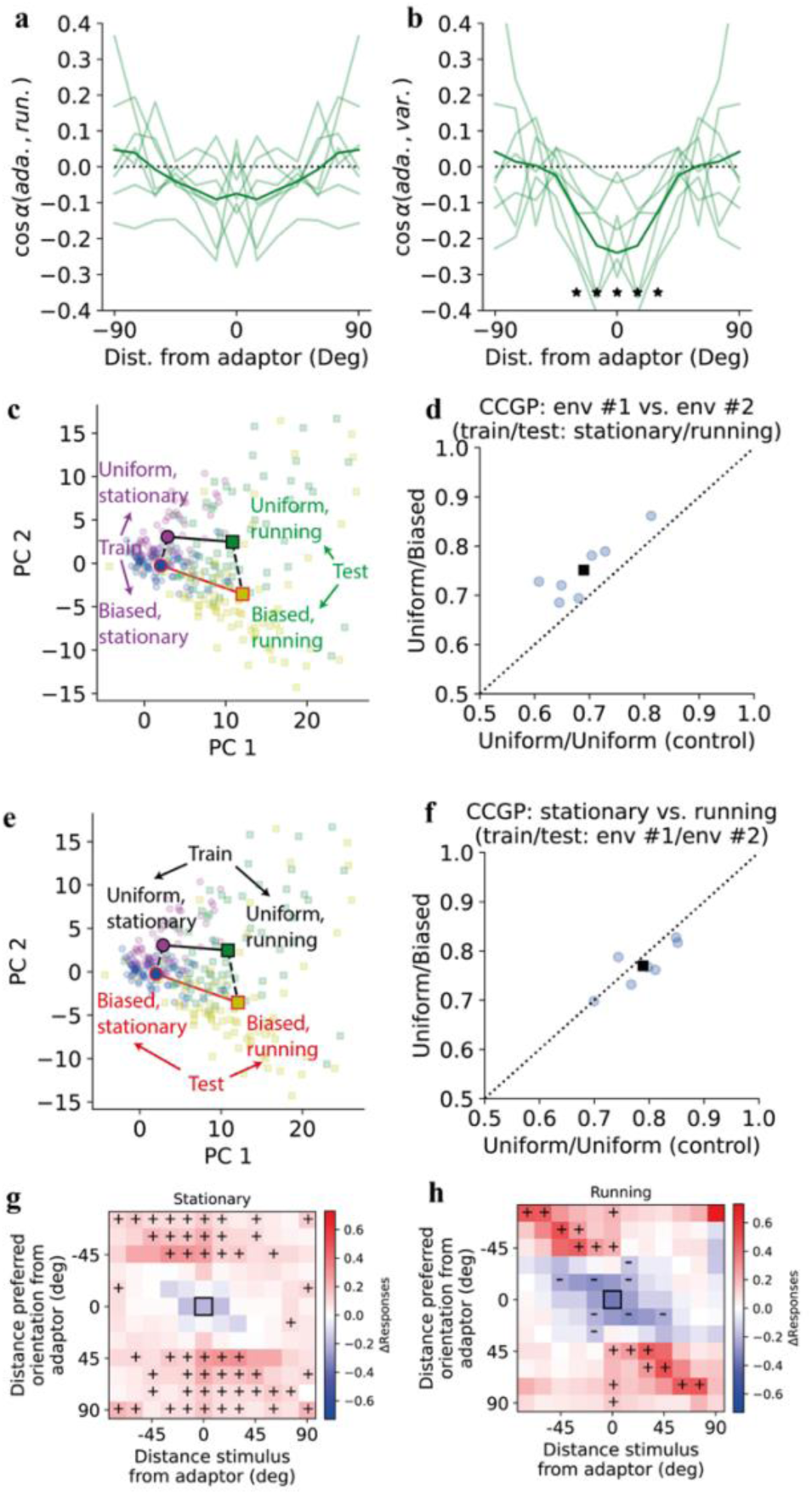
related to Figure 3. Adaptation-related changes align with the principal axis of response variability while remaining largely separable from locomotion. **a**) Angle between vector of adaptation changes and vector or locomotion changes; **b**) same as in (a) but the vector of locomotion changes is replaced by a vector aligned with the principal component of response trials for a given stimulus. Asterisks correspond to significant differences in angle from 0 deg (orthogonal vectors). **c**) responses to the same angle (adaptor at 45 deg) in different locomotion conditions (circles: stationary, squares: running) and adaptation conditions (magenta and green: uniform, blue and yellow: biased); the text is colored based on the analysis in (d), training and testing sets have also been inverted. **d**) Cross-condition generalization performance (CCGP) measured by discriminating a uniform block vs. another uniform block (x-axis, not shown in c) or a uniform block vs. a biased block (y-axis, shown in c) in stationary or running trials after training the model in the other condition locomotion condition. Circles are recording sessions with a sufficient number of stationary and running trials, and the black square is average across these conditions. **e**) Same as (c) but text is colored based on the analysis in f. **f**) Same as in (d) but CCGP is measured by discriminating stationary vs. running conditions in biased trials after training the model in a uniform block and testing it in another uniform block (x-axis, not shown in e) or training the model in a uniform block and testing it in a biased block. **g)** Difference in normalized responses between biased and uniform environments averaged across experiments and for the stationary condition only, similar to Fig. 3d, but for all cells rather than only selective cells. **h**) Similar to (g) but for the running condition.

**Supplementary Figure 13.**
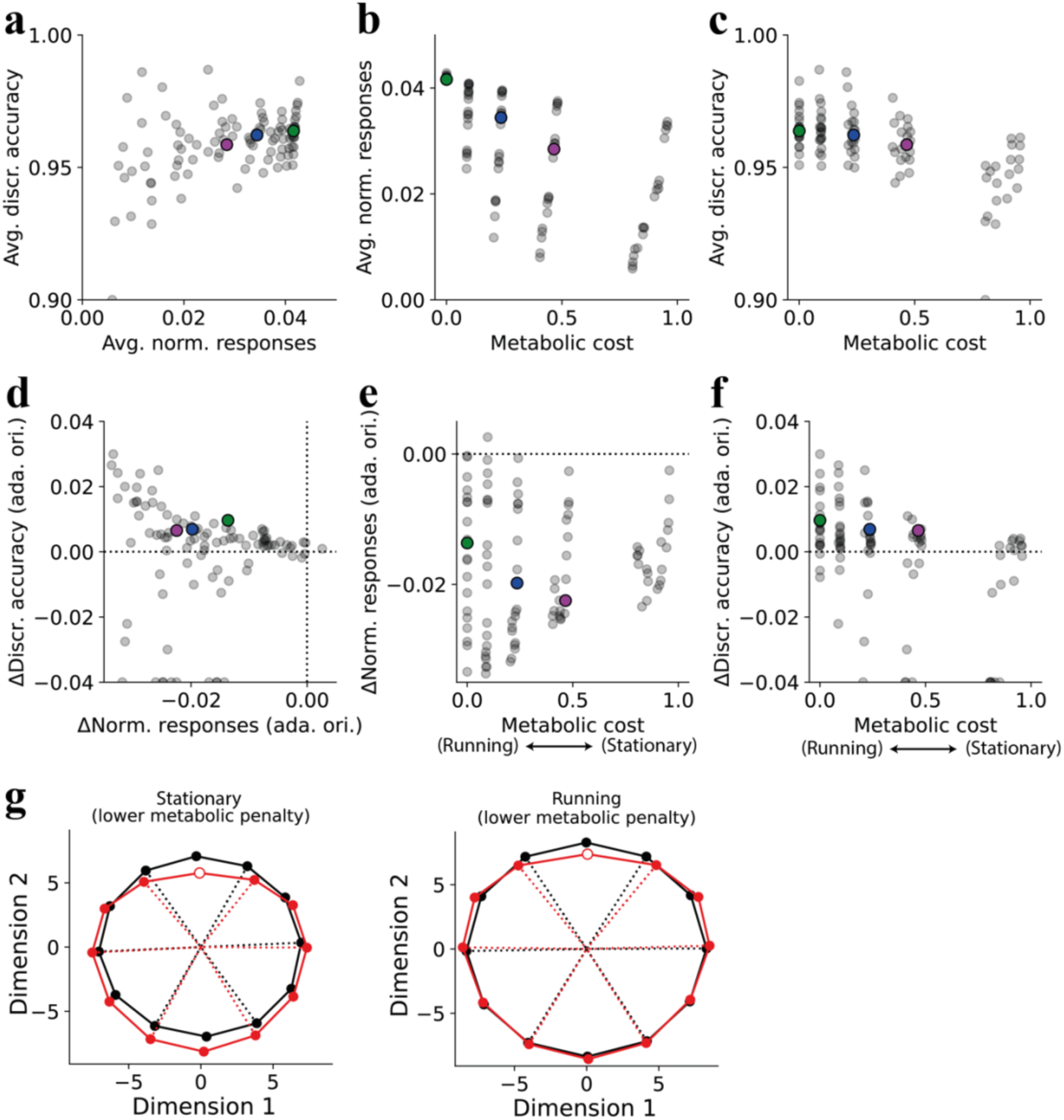
related to Figure 4. Varying the model’s metabolic cost reproduces the locomotion-like tradeoff between response magnitude and discriminability. **a**) Average discrimination accuracy (across pairs of orientations and environments) as a function of average response magnitude (across neurons, orientations, and environments) for different model’s hyperparameters; here and in the following panels, the magenta circle corresponds to an example (averaged across repeats of the same hyperparameter) stationary locomotion condition with metabolic penalty *λ* = 0.5, green circle is an example running condition with *λ* = 0; finally, the blue circle corresponds to an intermediate locomotion condition with *λ* = 0.25 and it is the example solution in Figure 4. In all these examples, we used the input noise level *ε*_*x*_ = 1 and number of hidden units *n*_*r*_ = 3,000. **b**) Average response magnitude as a function of metabolic cost. **c**) average discriminability as a function of metabolic cost. **d**) Changes in discrimination accuracy near the adaptor as a function of changes in responses for neurons tuned to the adaptor at the adaptor orientation for different models’ hyperparameters (same as in Figure 4h but with the example stationary and running conditions in purple and green, respectively). **e**) Changes in responses for neurons tuned to the adaptor at the adaptor orientation as a function of metabolic penalty. **f**) Changes in discrimination accuracy near the adaptor as a function of metabolic penalty. **g**) PCA computed independently per conditions, followed by a rigid alignment of orientations. Reducing metabolic costs during running leads to an expansion of distances in the high-dimensional spaces (visualized in low dimension) for one selected model.

**Supplementary Figure 14.**
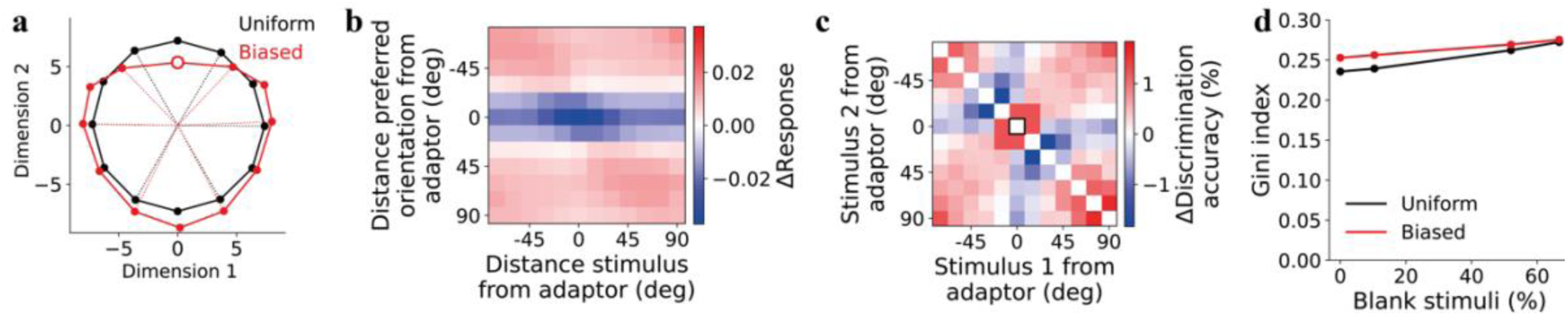
related to Figure 4. Adaptation effects are reproduced without blank stimuli, and sparsity is intrinsic to the model. All panels use the chosen Fig. 4 model (n_hidden_ = 3000, metabolic weight = 0.25, input noise = 1.0). **a**) Hidden-layer responses of one autoencoder trained with 0% blank stimuli, in the uniform (black) and biased (red) environments, projected and aligned as in Fig. 4c; adaptor marked (open red circle). **b**) Difference in superneuron responses between biased and uniform environments (as in Fig. 4d), for 0% blank stimuli (mean of 3 training repeats). **c**) Difference in discrimination accuracy between biased and uniform environments (as in Fig. 4f), for 0% blank stimuli (mean of 3 training repeats). **d**) Gini index across neurons, averaged over the 12 gratings (higher = sparser), versus the percentage of blank stimuli during training, for uniform (black) and biased (red) environments; total presentations held at 3000 (mean of 3 training repeats, s.d. < 0.001).

**Supplementary Figure 15.**
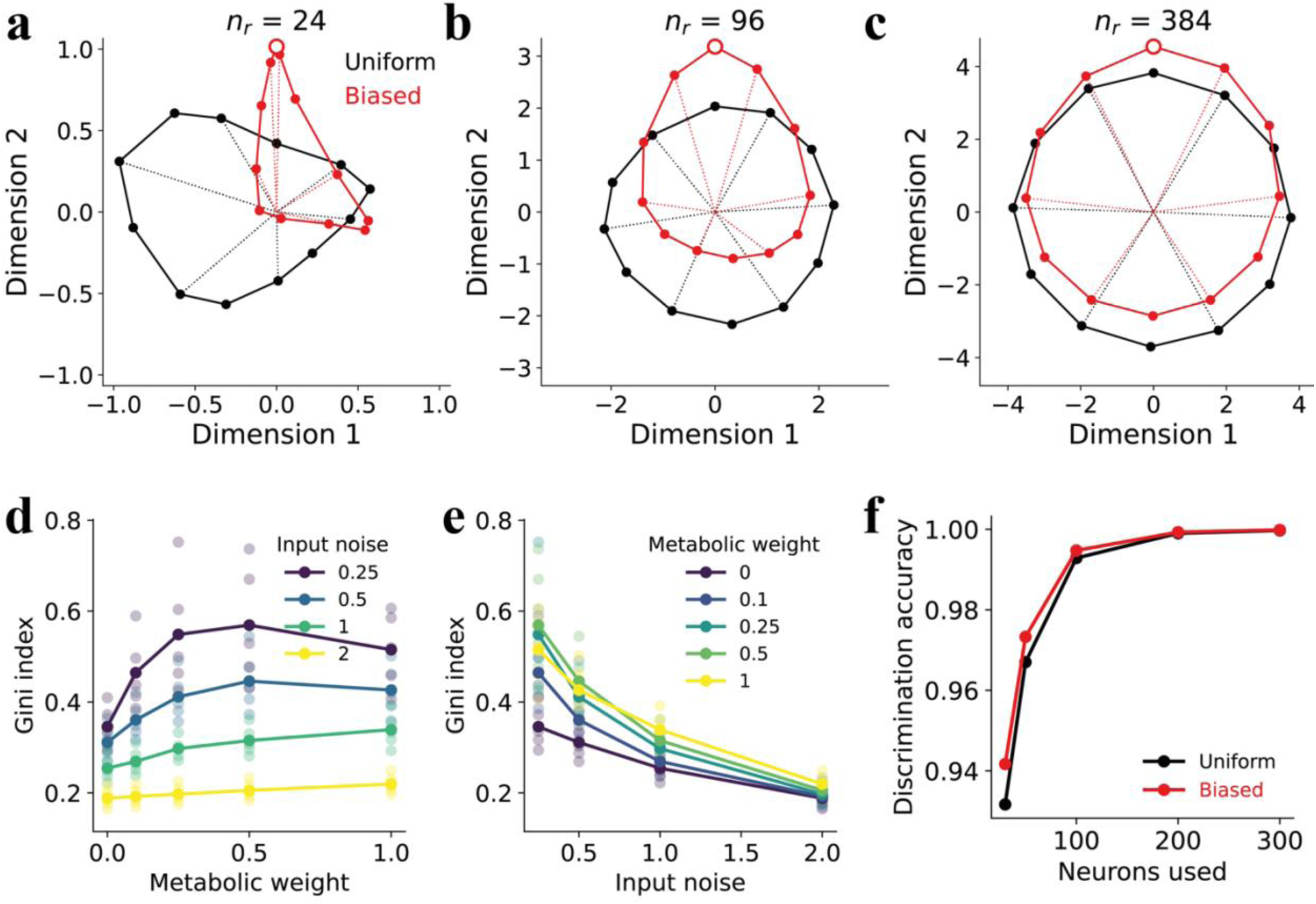
related to Figure 4. Reproducing the adaptation geometry requires a sufficient number of hidden units. **a**) Hidden-layer responses of one autoencoder with 24 hidden units, projected and aligned as in Fig. 4c**. b,c**) Same as (a) but for 96 and 384 hidden units. **d,e**) Gini index as a function of metabolic weights and input noise. A higher Gini index value indicates greater sparsity. **f**) Discrimination accuracy between two stimuli 15 degrees apart from the adaptor (30 degrees apart from each other) as a function of the number of hidden units (randomly sampled) used by the read-out unit.

**Supplementary Figure 16.**
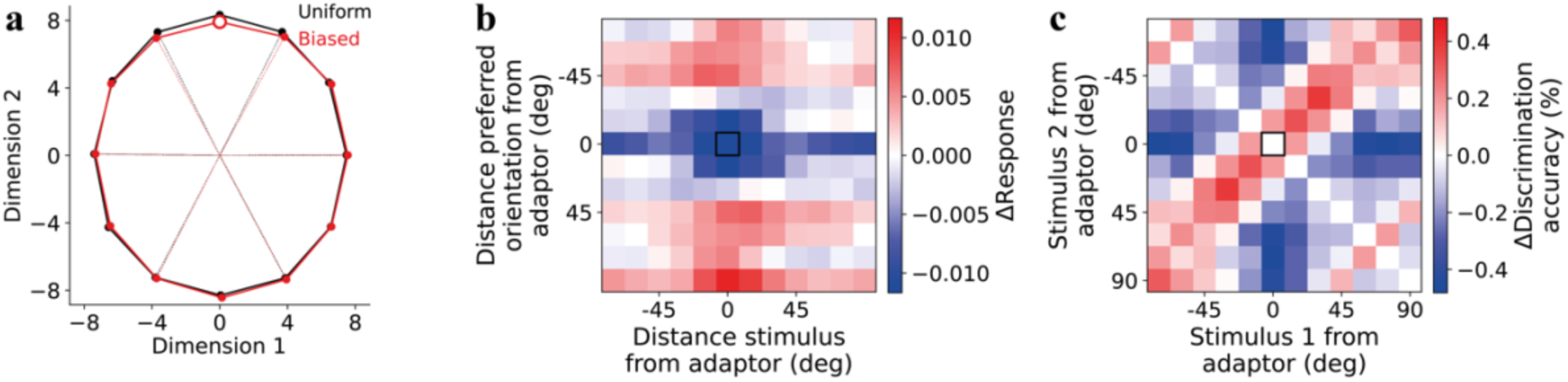
related to Figure 4. A convolutional autoencoder with far fewer parameters reproduces the adaptation-induced response suppression near the adaptor. **a**) Hidden-layer responses of one such autoencoder (metabolic weight = 0.25), in the uniform (black) and biased (red) environments, projected and aligned as in Fig. 4c; adaptor marked (open red circle). **b**) Difference in superneuron responses between biased and uniform environments (as in Fig. 4d), averaged across 30 autoencoders with the same architecture. **c**) Difference in discrimination accuracy between biased and uniform environments (as in Fig. 4f), averaged across the same 30 autoencoders.

**Supplementary Figure 17.**
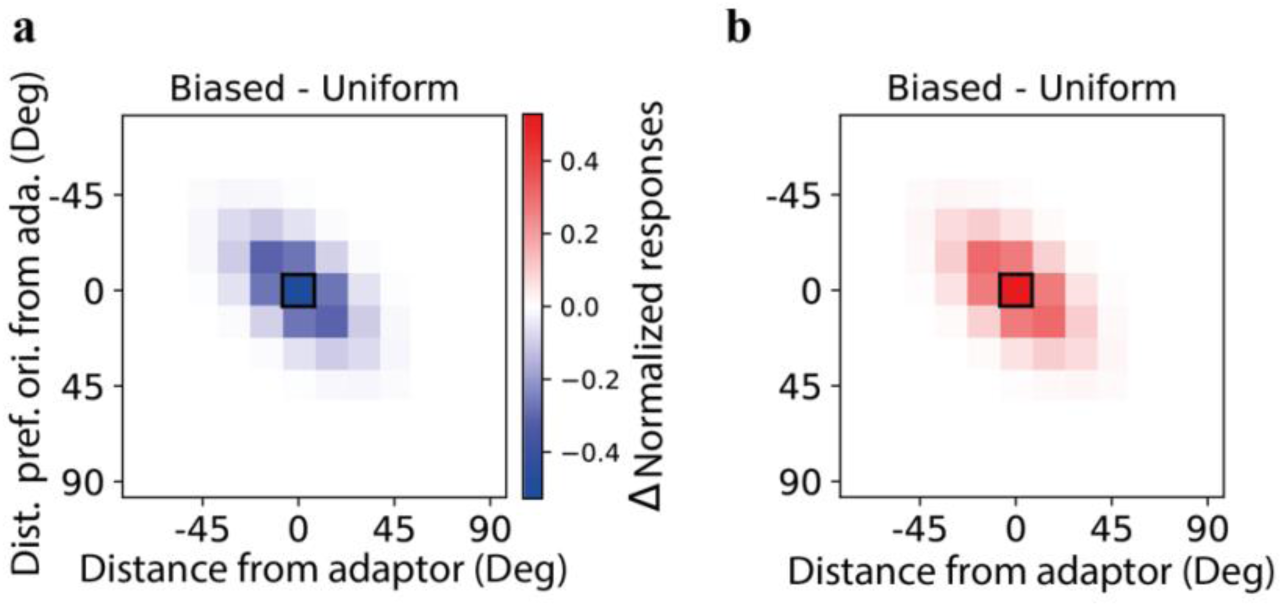
related to Figure 5. Simple localized response depression or facilitation. Differences in normalized responses of neurons binned by their preferred orientation between biased and uniform environments. The black square corresponds to the response of neurons tuned to the adapter to the adapter. **a,b**) Changes following tuning curves in Figure 5a,b.

